# Ketamine disinhibits dendrites and enhances calcium signals in prefrontal dendritic spines

**DOI:** 10.1101/659292

**Authors:** Farhan Ali, Danielle M. Gerhard, Katherine Sweasy, Santosh Pothula, Christopher Pittenger, Ronald S. Duman, Alex C. Kwan

## Abstract

A subanesthetic dose of ketamine causes acute psychotomimetic symptoms and then more sustained antidepressant effects. A key targeted brain region is the prefrontal cortex, and the prevailing disinhibition hypothesis posits that N-methyl-d-aspartate receptor (NMDAR) antagonists such as ketamine may act preferentially on GABAergic neurons. However, cortical GABAergic neurons are heterogeneous. In particular, somatostatin-expressing (SST) interneurons selectively inhibit dendrites and regulate synaptic inputs, yet their response to systemic NMDAR antagonism is unknown. Here, we report that administration of ketamine acutely suppresses the activity of SST interneurons in the medial prefrontal cortex of the awake mouse. The deficient dendritic inhibition leads to greater synaptically evoked calcium transients in the apical dendritic spines of pyramidal neurons. By manipulating NMDAR signaling via GluN2B knockdown, we show that ketamine’s actions on the dendritic inhibitory mechanism has ramifications for frontal cortex-dependent behaviors and cortico-cortical connectivity. Collectively, these results demonstrate dendritic disinhibition and elevated calcium levels in dendritic spines as important local-circuit alterations driven by the administration of subanesthetic ketamine.

## Introduction

Ketamine is a non-competitive N-methyl-d-aspartate receptor (NMDAR) antagonist. In humans, systemic administration of ketamine at a subanesthetic dose have both acute and sustained effects. Acutely, within about 1 hour following administration, subjects experience psychotomimetic symptoms, perceptual aberrations, and cognitive impairments^1^. Because the acute effects mimic the positive and negative symptoms of schizophrenia, ketamine has found use as a tool for studying glutamatergic dysfunction and evaluating antipsychotic compounds^2, 3^. On a longer timescale, starting from 2 to 4 hours following administration, patients suffering from major depressive episodes report a robust antidepressant effect that remain for about a week^4, 5^. Although these behavioral effects are well established and have substantial translational implications, the mechanisms by which ketamine impacts neural activity remains unclear.

In rodents, intraperitoneal or subcutaneous injection of subanesthetic ketamine activates multiple, distributed brain regions, as indicated by brain-wide mapping of metabolic activity^6, 7^. Among the regions with elevated metabolic activity is the medial prefrontal cortex, including the ventral prelimbic and infralimbic sub-regions as well as the dorsal cingulate and secondary motor sub-regions (Cg1/M2). This locus is consistent with earlier works demonstrating that, for NMDAR antagonists, there is a dose-dependent increase in the extracellular levels of glutamate in the rat prefrontal cortex^8^ as well as an increase in the overall firing rate of prefrontal cortical neurons^9^. Since NMDAR is an ionotropic glutamate receptor that contributes to postsynaptic depolarization, why would an antagonist lead to more spiking activity? One prevailing idea is the disinhibition hypothesis, which posits that NMDAR antagonists at low dose may act preferentially on GABAergic neurons, thereby disinhibiting excitatory neurons^10, 11^. In support of the disinhibition hypothesis, fast-spiking GABAergic neurons, identified based on their narrow spike waveforms, exhibit a concomitant reduction in firing *in vivo* at the time of prefrontal hyperactivity during NMDAR hypofunction^12^.

However, GABAergic neurons are not homogeneous. Fast-spiking cells constitute a fraction of the GABAergic neurons, and another non-overlapping subpopulation is the somatostatin-expressing (SST) interneurons. SST interneurons are a major subpopulation that constitutes 20% of the GABAergic neurons in the mouse frontal cortex^13^. Unlike the fast-spiking cells which inhibit cell bodies, SST interneurons include the so-called Martinotti cells that selectively target apical dendrites and thus are poised to control synaptic inputs and dendritic integration^14^. The cell-type-specific differences suggest that if SST interneurons were altered, then the resulting circuit-level and behavioral impacts may be quite distinct from those involving perturbations of fast-spiking cells. Still unknown, however, is whether and how SST interneurons and dendritic inhibition may be impacted by subanesthetic ketamine.

In this study, we used two-photon calcium imaging to characterize SST interneurons, SST axons, pyramidal neurons, and pyramidal dendritic spines in awake, head-fixed mice. For each cellular compartment, we compared spontaneous calcium transients before and within an hour after the administration of subanesthetic ketamine, relative to saline control. We found that ketamine suppresses the activity of SST interneurons in the medial prefrontal cortex. We further show that this loss of dendritic inhibition contributes to elevated synaptic calcium transients in the apical dendritic spines of pyramidal neurons. Based on these results, we conclude that dendritic disinhibition and calcium elevations in dendritic spines are important components of ketamine’s actions on the prefrontal cortex.

## Results

### Subanesthetic ketamine modifies prefrontal cortical activity in a cell-type-specific manner

We performed two-photon microscopy on awake, head-fixed mice (**Fig. 1a**), targeting the Cg1/M2 sub-regions of the medial prefrontal cortex (**Fig. 1b**). Initially, while mice were head-fixed under the two-photon microscope, we recorded body motion using an infrared camera. This was because systemic administration of subanesthetic ketamine induces hyperlocomotion in rodents^15^, and we wanted to avoid movement as a confound in our imaging experiments. We observed that ketamine (10 mg/kg, s.c.) increased body motion but only transiently, and therefore limited all of our data collection to 30 – 60 minutes post-injection (**Fig. 1c, d**). For calcium imaging, we used AAV1-CamKII-GCaMP6f-WPRE-SV40 to express the calcium-sensitive fluorescent protein GCaMP6f in pyramidal neurons in Cg1/M2, and imaged spontaneous fluorescence transients from the awake mouse (**Fig. 1e**). From the fluorescence transients, we detected calcium events using a peeling method based on template matching^16^. Ketamine (10 mg/kg, s.c.) increased the rate of spontaneous calcium events in pyramidal neuron cell bodies in layer 2/3 (200 – 400 µm from the dura) (ketamine: 23.7 ± 2.1%, saline: 9.4 ± 1.9%, relative to pre-injection, mean ± s.e.m.; *P* = 3 × 10^-8^, two-sample t-test; **Fig. 1f, g; Supplementary Fig. 1a**). The elevation in calcium event rates did not correlate with animal movement (**Supplementary Fig. 1b**) and was consistent across individual mice (**Supplementary Fig. 2a**). Somatic calcium transients have been shown to directly relate to the firing rate of cortical neurons^17, 18^, therefore the elevated calcium event rates reflect hyperactivity of pyramidal neurons, consistent with previous reports^12, 19^. To characterize the effect of ketamine on the SST subpopulation of GABAergic neurons, we injected AAV1-Syn-DIO-GCaMP6s-WPRE-SV40 into Cg1/M2 of SST-IRES-Cre mice to express GCaMP6s selectively in SST interneurons (**Fig. 1h**). Following ketamine, the rate of spontaneous calcium events was reduced for the cell bodies of layer 2/3 SST interneurons (ketamine: -13 ± 3%, saline: 13 ± 6%, relative to pre-injection, mean ± s.e.m.; *P* = 1 × 10^-4^, two-sample t-test; **Fig. 1i, j, Supplementary Fig. 1c, d**; **Supplementary Fig. 2b**). These results indicate that ketamine increases the activity of pyramidal neurons while suppressing the activity of SST GABAergic neurons.

**Fig. 1.**
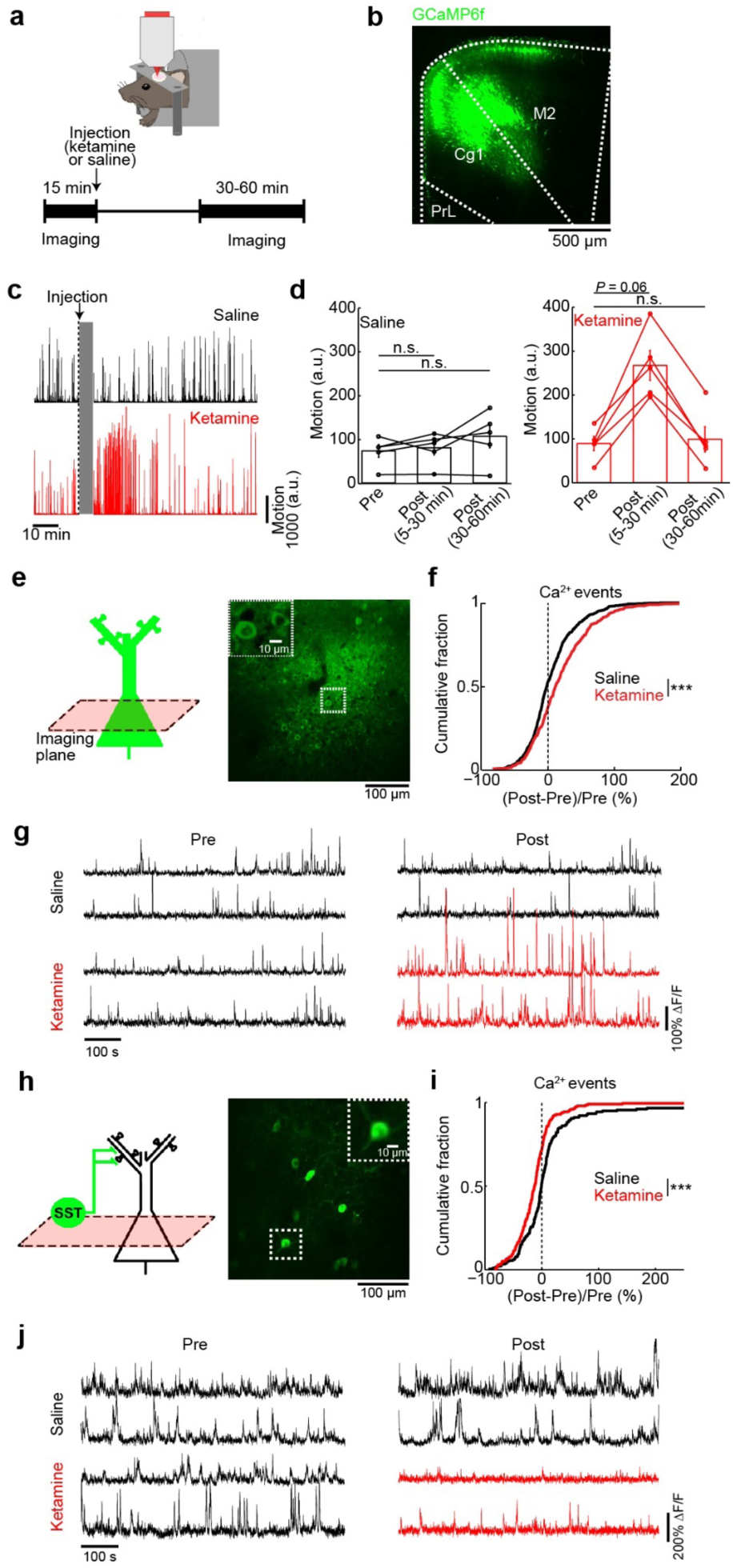
Effects of subanesthetic ketamine on the activity of pyramidal neurons and SST interneurons *in vivo*. (a) Schematic and timeline of the experiments. (b) Coronal histological section, showing the extent of AAV-mediated expression of GCaMP6f in the mouse medial prefrontal cortex. Cg1, cingulate cortex. M2, secondary motor cortex. PrL, prelimbic cortex. (c) Example motion traces from a head-fixed animal during a two-photon imaging session before and after saline injection (black), and another session before and after ketamine injection (red). Gray shading denotes the 5 minutes during which the injection was made and imaging was done to verify that the field of view had not shifted. (d) Time-averaged motion (mean ± s.e.m.) for epochs including pre-injection, 5 - 30 min post-injection, and 30 - 60 min post-injection for saline (left) and ketamine (right). For saline injection (black), there were no detectable differences between epochs (pre-injection versus 5-30 min post-injection: *P* = 0.6; pre-injection versus 30 - 60 min post-injection: *P* = 0.1; Wilcoxon signed rank test). For ketamine (red), hyperlocomotion was detected transiently following injection (pre-injection versus 5-30 min post-injection: *P* = 0.06; pre-injection versus 30 - 60 min post-injection: *P* = 1; Wilcoxon signed rank test). Each point is an imaging session. n = 5 animals each for saline and ketamine. (e) Schematic of imaging location, and an *in vivo* two-photon image of GCaMP6s-expressing pyramidal neurons in Cg1/M2. Inset, magnified view of neuronal cell bodies (f) The normalized difference in the rate of spontaneous calcium events of pyramidal neurons. Normalized difference was calculated as post-minus pre-injection values normalized by the pre-injection value (ketamine: 23.7 ± 2.1%, saline: 9.4 ± 1.9%, mean ± s.e.m.; *P* = 3 × 10^-8^, two-sample t-test). For ketamine, *n* = 613 cells from 5 animals. For saline, *n* = 681 cells from 5 animals. (g) Each row shows time-lapse fluorescence transients from the same pyramidal cell in the pre-(left) and post-injection (right) periods. Two example spines were plotted for saline injection (black), and two other examples were plotted for ketamine injection (pre-injection: black; post-injection: red). (h - j) Same as (e-g) for GCaMP6s-expressing SST interneurons in Cg1/M2 of SST-IRES-Cre animals (ketamine: -12 ± 3%, saline: 13 ± 6%, mean ± s.e.m.; *P* = 1 x× 10^-4^, two-sample t-test). For ketamine, *n* = 198 cells from 5 animals. For saline, *n* = 179 cells from 5 animals. * *P* < 0.05; ** *P* < 0.01; *** *P* < 0.001; n.s., not significant

**Fig. 2.**
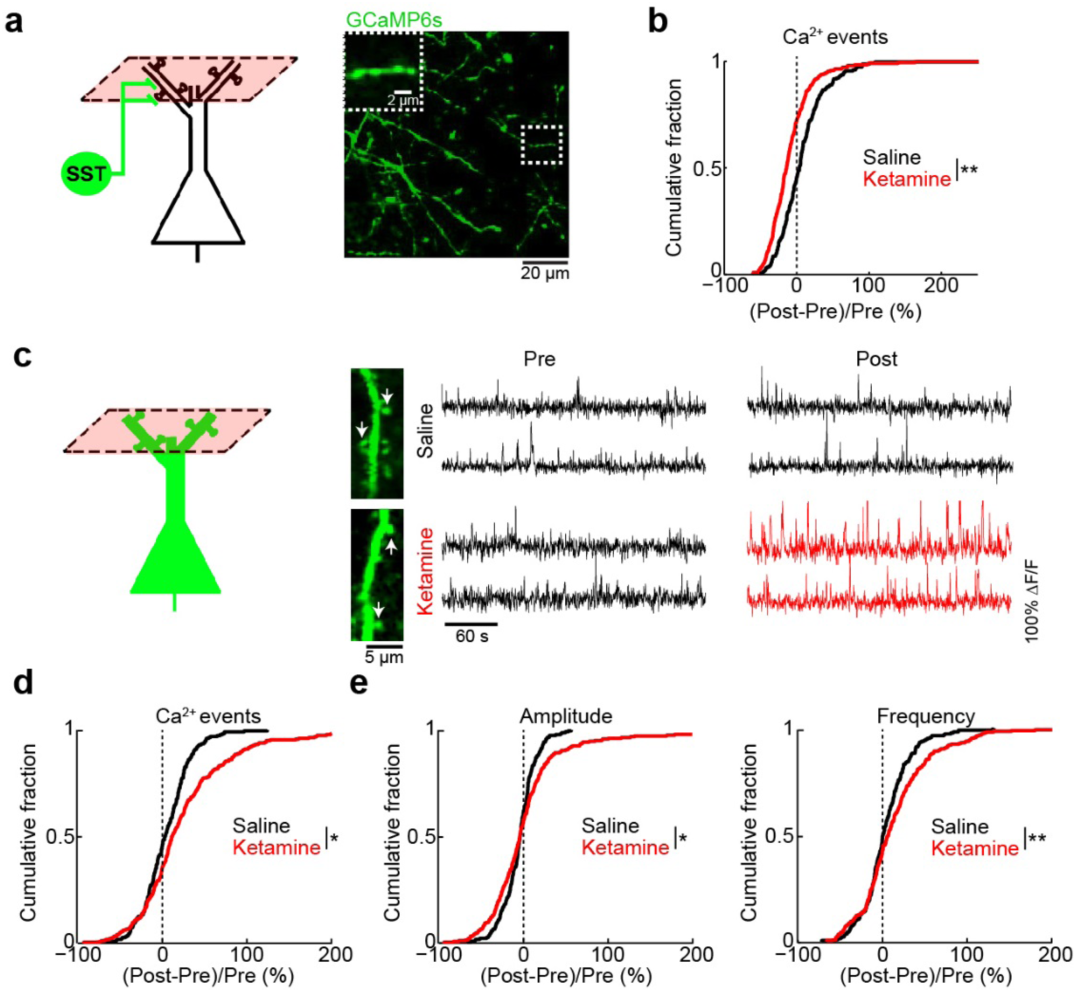
Ketamine induces opposing effects on dendrite-targeting SST axons and apical dendrites of pyramidal neurons. (a) Schematic of imaging location, and an *in vivo* two-photon image of GCaMP6s-expressing SST axons in superficial layers of Cg1/M2 of SST-IRES-Cre animals. Inset, magnified view of a SST axonal segment. (b) The normalized difference in the rate of spontaneous calcium events for SST axons in superficial layers. Normalized difference was calculated as post-minus pre-injection values normalized by the pre-injection value (ketamine: -9 ± 3%, saline: 12 ± 7%, mean ± s.e.m.; *P* = 0.002, two-sample t-test). For ketamine, *n* = 269 boutons from 5 animals. For saline, *n* = 214 boutons from 5 animals. (c) Left, schematic of imaging location. Right, each row shows time-lapse fluorescence transients from the same dendritic spine in the pre-(left) and post-injection (right) periods. Two example spines were plotted for saline injection (black), and two other examples were plotted for ketamine injection (pre-injection: black; post-injection: red). Locations of the spines in the *in vivo* two-photon images are indicated by white arrows (d) Same as (b) but for dendritic spines in superficial layers of Cg1/M2 (ketamine: 43.42 ± 0.01%, saline: 4.34 ± 0.01%, mean ± s.e.m.; *P* = 0.02, two-sample t-test). For ketamine, *n* = 280 dendritic spines from 5 animals. For saline, *n* = 231 dendritic spines from 5 animals. (e) The normalized difference in amplitude (ketamine: 5 ± 4%, saline: -4 ± 1%, mean ± s.e.m.; *P* = 0.03, two-sample t-test), and frequency of binned calcium events (ketamine: 16 ± 4%; saline: 4 ± 2%; *P* = 0.008, two-sample t-test). * *P* < 0.05; ** *P* < 0.01; *** *P* < 0.001; n.s., not significant.

### Ketamine disinhibits dendrites and increases calcium transients in dendritic spines

SST interneurons inhibit selectively dendrites including the dendritic spines of pyramidal neurons ^14^. The ketamine-induced reduction of SST interneuron activity is therefore expected to diminish dendritic inhibition. To determine if this may apply to the apical dendritic compartment, we imaged the inhibitory inputs – SST axonal boutons in the superficial layer of Cg1/M2 (<200 µm from the dura) (**Fig. 2a**), which showed fewer calcium events after ketamine (**Fig. 2b; Supplementary Fig. 2c**). We also imaged directly the postsynaptic calcium signals in the apical dendrite tufts of pyramidal neurons. We focused on the dendritic spines (**Fig. 2c**), because subthreshold synaptic activation can be characterized by their accompanying calcium elevations in the spine compartment with minimal outflow to the dendritic shaft ^20^. To restrict our analyses to these localized fluorescence transients, we used a standard procedure to regress out the contribution from the dendritic shaft^18, 20^, following which we applied the same calcium event detection procedure. Ketamine induced a 43.42 ± 0.01% increase in spontaneous calcium events for dendritic spines in Cg1/M2, significantly higher than saline controls (4.34 ± 0.01%, *P* = 0.02, two-sample t-test; **Fig. 2c, d; Supplementary Fig. 2d**). To characterize the temporal dynamics of the aberrant transients, we identified calcium events that co-occurred in each image frame, thereby separating amplitude (number of events in each frame) from frequency (number of frames with at least one event). Both the amplitude and frequency parameters were elevated following ketamine (**Fig. 2e**). The results did not depend on the specifics of the template in the event detection procedure, because an alternative analysis based on threshold crossing yielded similar conclusions. (**Supplementary Fig. 3**). Interestingly, the ketamine-induced elevation of spine calcium transients was regionally specific, because there was little effect when we repeated the experiment in the primary motor cortex (**Fig. 3; Supplementary Fig. 2e**). Altogether, the results demonstrate that ketamine disinhibits apical dendrites, leading to elevated calcium transients in the apical dendritic spines of prefrontal pyramidal neurons.

**Fig. 3.**
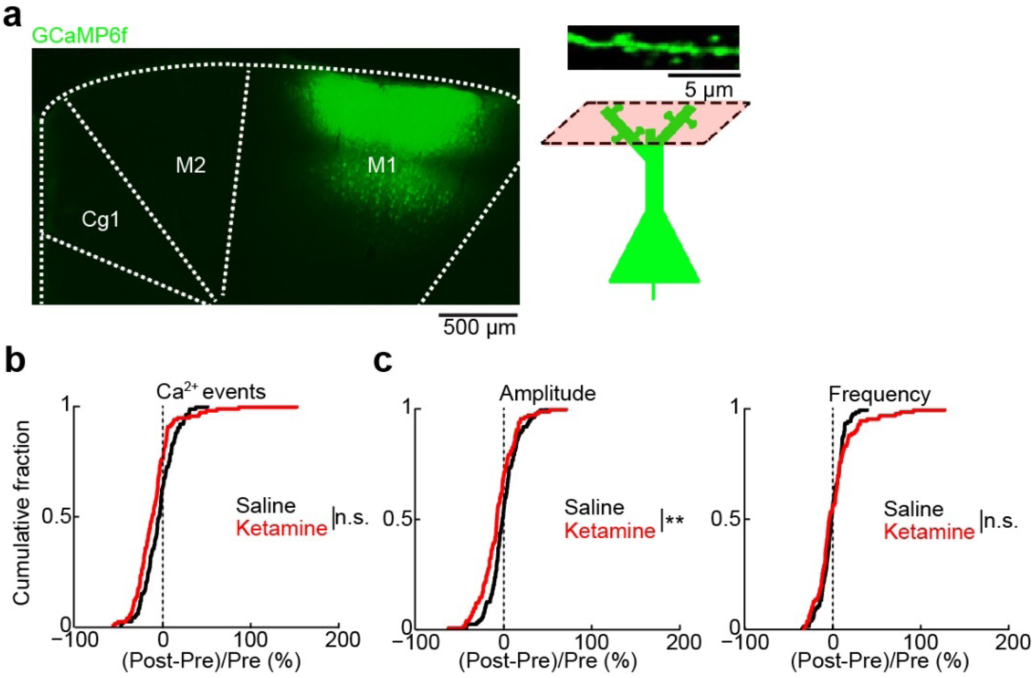
Effect of ketamine on calcium dynamics for dendritic spines in the primary motor cortex. (a) Left, coronal histological section, showing the extent of AAV-mediated expression of GCaMP6f in the primary motor cortex, M1. Right, schematic of imaging location. (b) The normalized difference in the rate of spontaneous calcium events for apical dendritic spines in M1. Normalized difference was calculated as post-minus pre-injection values normalized by the pre-injection value (ketamine: -7 ± 2%, mean ± s.e.m.; saline: -2 ± 2% for saline; *P* = 0.05, two-sample t-test). For ketamine, *n* = 124 dendritic spines from 3 animals. For saline, *n* = 120 dendritic spines from 3 animals. (c) The normalized difference in amplitude (ketamine: -7 ± 2%; saline: -5 ± 2%; *P* = 0.004, two-sample t-test) and frequency of binned calcium events (2 ± 2% for ketamine; -1 ± 1% for saline; *P* = 0.3, two-sample t-test). * *P* < 0.05; ** *P* < 0.01; *** *P* < 0.001; n.s., not significant.

### Elevated synaptic calcium responses to long-range cortical inputs

By focusing on fluorescence transients that are localized to dendritic spines, we are characterizing calcium signals related to synaptic activation. It follows then that a more stringent and explicit test for a synaptically activated signal is to directly stimulate the afferent inputs. To test this, we take advantage of the knowledge that apical dendritic spines receive long-range cortico-cortical inputs^21^, and Cg1/M2 receives an abundance of monosynaptic excitatory inputs from the retrosplenial cortex (RSC). We implanted electrodes to stimulate RSC (biphasic pulse with duration of 10 ms and a peak amplitude of ±150 µA), and simultaneously imaged the evoked calcium responses in Cg1/M2 (**Fig. 4a**). First, we confirmed that electrical stimulation activated the long-range inputs, by imaging the axons emanating from RSC in Cg1/M2 and finding stimulation-aligned calcium transients (**Supplementary Fig. 4a-e**). Then, in a separate group of animals, we imaged the evoked calcium responses in single dendritic spines in Cg1/M2 while varying the stimulation strength (number of pulses = 1, 2, 4, 8, 16, 32, or 64 per trial; **Fig. 4b**). Ketamine, but not saline, increased evoked responses in Cg1/M2 dendritic spines (ketamine: F(6,1146) = 6.6, *P =* 7 × 10^-7^, saline: F(6,1218) = 20.0, *P* = 0.06, two-way interaction between time (pre-vs. post) and stimulation strength, two-way mixed ANOVA after three-way mixed ANOVA; **Fig. 4c, d**). Across all imaged spines, the increase in evoked calcium response was more pronounced at higher stimulation strengths (**Fig. 4d; Supplementary Fig. 4f-g**). By directly controlling the activation of long-range inputs, these results provide evidence that ketamine elevates the synaptically evoked component of calcium transients in prefrontal dendritic spines.

**Fig. 4.**
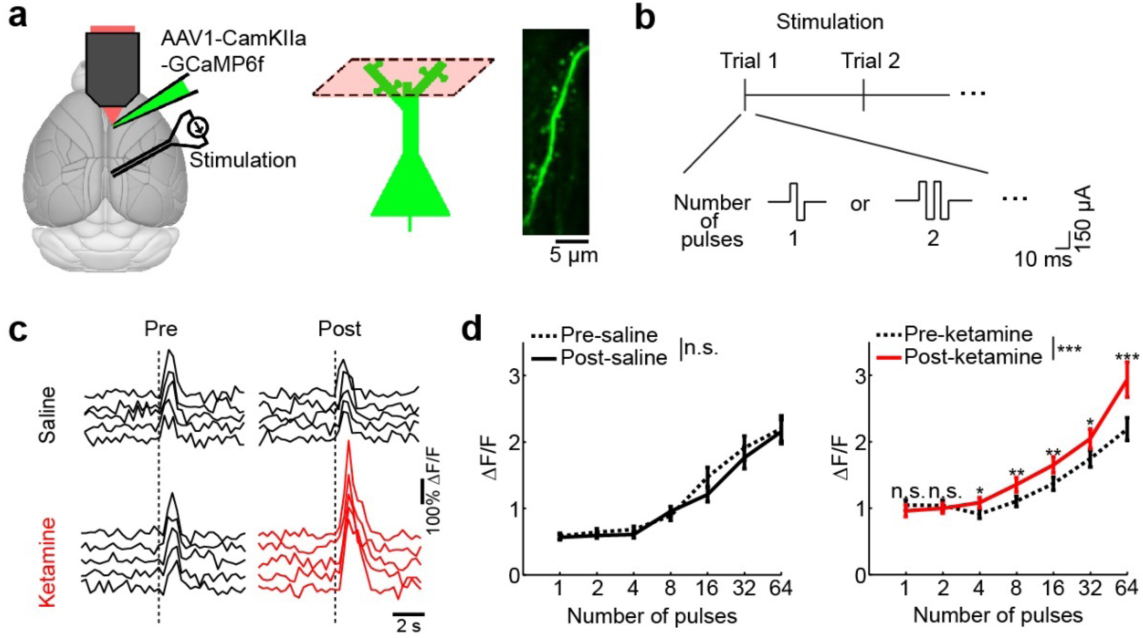
Ketamine increases synaptically evoked calcium responses in dendritic spines. (a) Schematic of experimental setup and imaging location. Right, an *in vivo* two-photon image of GCaMP6f-expressing apical dendritic spines in the superficial layer of Cg1/M2. (b) Protocol for the RSC stimulation. (c) Fluorescence transients in dendritic spines in Cg1/M2 in response to electrical stimulation of the retrosplenial cortex (RSC). Each trace shows a single 32-pulse-stimulation trial during pre-(left) or post-injection (right) period. One example spine was plotted for saline injection (black), and another example spine was plotted for ketamine injection (red). Dashed line, time of stimulation onset. (d) Trial-averaged spine calcium responses as a function of the number of stimulation pulses applied in a trial, for pre-(dashed line) versus post-injection period (solid line) and for saline (black line) versus ketamine injection (red line). A three-way mixed ANOVA was performed with drug (saline, ketamine) as a between-subjects factor, and stimulation levels (1, 2, 4, 8, 16, 32, 64) and epoch (pre-, post-injection) as within-subjects factors. A three-way interaction was significant (F(6,2364) = 5.6, *P* = 9 × 10^-6^). A subsequent two-way mixed ANOVA was performed separately on the saline and ketamine datasets, which showed a significant two-way interaction between time (pre-vs. post) and stimulation levels for ketamine (F(6,1146) = 6.6, *P =* 7 × 10^-7^), but not for saline (F(6,1218) = 20.0, *P* = 0.06). Post-hoc Tukey-Kramer test on the ketamine dataset indicated significantly elevated spine responses for post-vs. pre-injection for stimulation pulse numbers of 4 (*P* = 0.01), 8 (*P* = 0.007), 16 (*P* = 0.008), 32 (*P =* 0.02), and 64 (*P* = 9 × 10^-5^). Line, mean ± s.e.m. For saline, *n* = 204 dendritic spines from 4 animals. For ketamine, *n* = 192 dendritic spines from 4 animals. * *P* < 0.05; ** *P* < 0.01; *** *P* < 0.001; n.s., not significant. Error bars, ± s.e.m.

### GluN2B expression in SST interneurons mediates the effect of ketamine on dendritic calcium signals

We wanted to determine the causal relations between ketamine, SST interneuron activity, and synaptic calcium signals. We predict that if ketamine exerts its effects through SST interneurons, then knocking down NMDAR signaling in SST interneurons should reproduce the local-circuit alterations and render additional ketamine ineffective. To test this prediction, we created AAV1-CMV-dsRed-pSico-GluN2BshRNA to induce Cre-dependent expression of short hairpin RNA (shRNA) against GluN2B (**Fig. 5a**)^22, 23^. We targeted the GluN2B subunit, because neocortical SST interneurons contain abundant GluN2B transcripts^24, 25^, and the subunit contributes a sizable fraction of NMDAR-mediated current in the related low-threshold spiking interneurons in the rodent medial prefrontal cortex^26^. This virus was confirmed to reduce GluN2B expression in a Cre-dependent manner *in vitro* (−42 ± 3% relative to controls; **Supplementary Fig. 5**) and decrease NMDAR-mediated currents in brain slices (Gerhard et al., manuscript in preparation). Injected into Cg1/M2 of SST-IRES-Cre animals (**Fig. 5b**), viral-mediated knockdown of GluN2B in SST interneurons (GluN2B-SST KD) reduced SST interneuron activity at baseline (GluN2B-SST KD: 1.5 ± 0.2 Hz, controls: 1.9 ± 0.1 Hz, mean ± s.e.m.; *P* = 0.04, two-sample t-test; **Fig. 5c**), mimicking the effects of ketamine. Notably, GluN2B-SST KD blocked further ketamine-induced influence on SST interneuron activity (**Fig. 5d**). In a separate group of animals, we imaged dendritic spines, finding that GluN2B-SST KD increased the rate of spontaneous calcium events at baseline (GluN2B-SST KD: 0.85 ± 0.03 Hz, controls: 0.73 ± 0.04 Hz; *P* = 0.01, two-sample t-test; **Fig. 5e**), and additional ketamine had no detectable effect (**Fig. 5f**). The causal manipulations thus confirm the predicted role of SST interneurons and highlight the involvement of GluN2B in mediating ketamine’s effect on the dendritic spines.

**Fig. 5.**
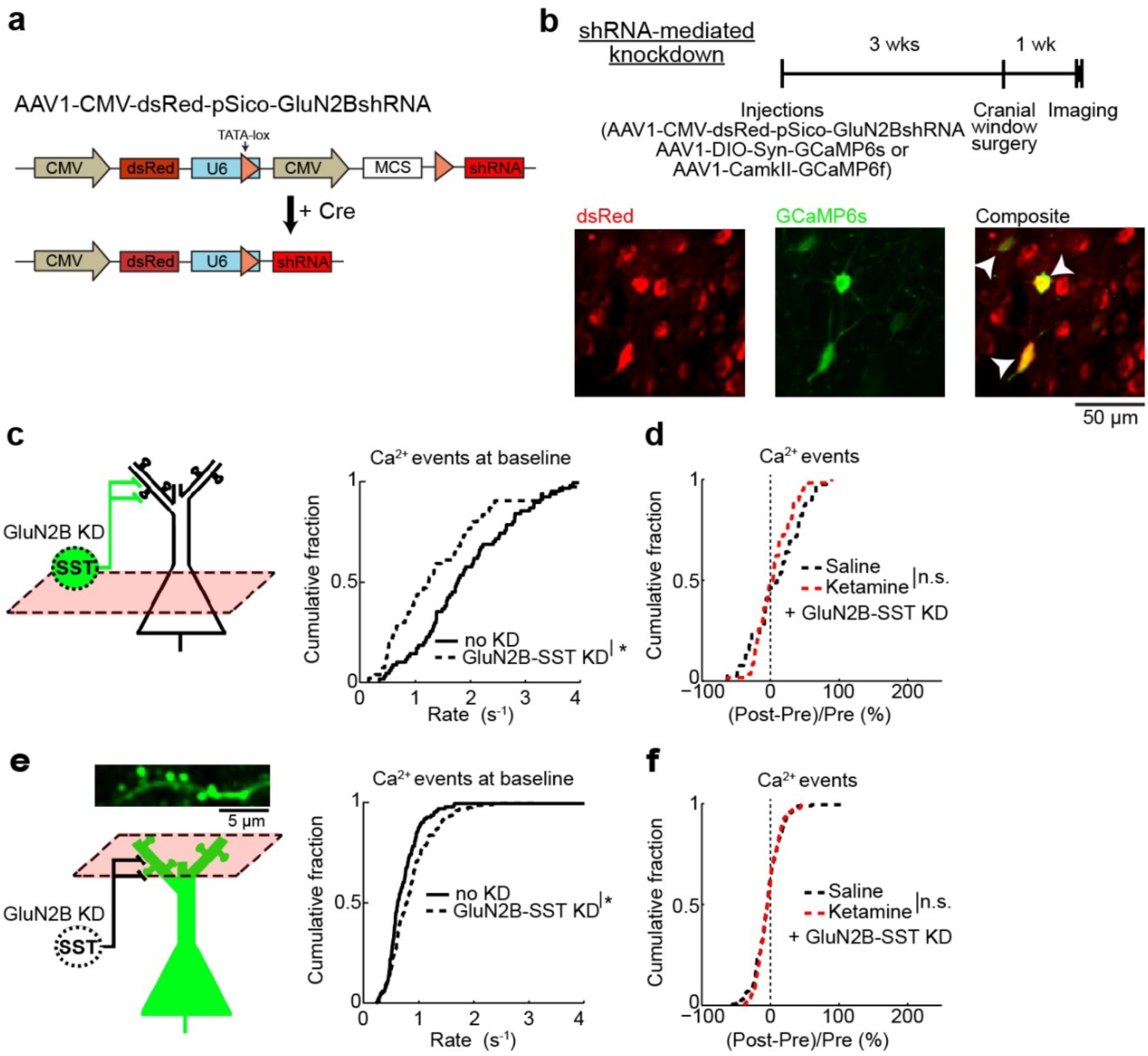
GluN2B expression in SST interneurons mediates effect of ketamine on dendritic spine calcium. (a) Schematic of AAV1-CMV-dsRed-pSico-GluN2BshRNA (standard AAV cassettes not drawn) for Cre-dependent expression of shRNA against GluN2B. (b) Top, timeline for experiments involving selective knockdown of GluN2B expression in SST interneurons (GluN2B-SST KD). Bottom, *in vivo* two-photon image of Cre-independent dsRed expression (left), Cre-dependent GCaMP6s expression (middle), and a composite image (right). Arrowheads, cells that expressed both dsRed and GCaMP6s. (c) The rate of spontaneous calcium events for GCaMP6s-expressing SST interneuron cell bodies with or without GluN2B-SST KD (GluN2B-SST KD: 1.5 ± 0.2 Hz, control SST-IRES-Cre animals with no KD: 1.9 ± 0.1 Hz, mean ± s.e.m.; *P* = 0.04, two-sample t-test). For GluN2B-SST KD, *n* = 58 cells from 3 animals. For control animals, *n* = 72 cells from 3 animals. (d) The normalized difference in the number of spontaneous calcium events GCaMP6s-expressing SST interneuron cell bodies, for GluN2B-SST KD with ketamine or saline injection. Normalized difference was calculated as post-minus pre-injection values normalized by the pre-injection value (ketamine with GluN2B-SST KD: 2 ± 4%, saline with GluN2B-SST KD: 7 ± 6%, mean ± s.e.m.; *P* = 0.46, two-sample t-test). For ketamine, *n* = 58 cells from 3 animals. For saline, *n* = 39 cells from 3 animals. (e) Same as (c) for GCaMP6f-expressing apical dendritic spines (GluN2B-SST KD: 0.85 ± 0.03 Hz, mean ± s.e.m., control SST-IRES-Cre animals: 0.73 ± 0.04 Hz, mean ± s.e.m.; *P* = 0.01, two-sample t-test). For GluN2B-SST KD, *n* = 277 spines from 4 animals. For control animals, *n* = 118 spines from 3 animals. (f) Same as (d) for GCaMP6f-expressing apical dendritic spines (ketamine with GluN2B-SST KD: -4 ± 1%, saline with GluN2B-SST KD: -5 ± 2%, mean ± s.e.m.; *P* = 0.8, two-sample t-test). For ketamine, *n* = 169 spines from 4 animals. For saline, *n* = 171 spines from 4 animals. * *P* < 0.05; ** *P* < 0.01; *** *P* < 0.001; n.s., not significant.

### Effects of ketamine on a subset of behaviors depend on dendritic inhibition

Ketamine exerts numerous acute behavioral effects, so next we asked to what extent dendritic disinhibition in the medial prefrontal cortex may be involved. We tested animals on three behavioral assays: trace fear conditioning, pre-pulse inhibition, and locomotor activity, and determined the effect of ketamine with or without bilateral GluN2B-SST KD (**Fig. 6a**). In trace fear conditioning, an auditory stimulus is paired with a footshock with an intervening trace period. This assay requires associative learning across a temporal gap, which relies on the medial prefrontal cortex^27^. SST-IRES-Cre animals received ketamine (10 mg/kg, s.c.) 15 minutes before conditioning, and subsequently showed impaired freezing 24 hours later in the 15-s trace but not in the delay (no trace) paradigm (**Fig. 6b**), indicating an impairment specific to temporal association but not learning in general. After bilateral GluN2B-SST KD in Cg1/M2, ketamine no longer had detectable effects on trace fear learning (**Fig. 6b**). Pre-pulse inhibition, a cross-species measure of sensorimotor gating, is regulated by prefrontal cortical activity^28^. Ketamine reduced pre-pulse inhibition in control SST-IRES-Cre animals (main effects of pre-pulse intensity, F(2,12) = 27.5, *P* = 4 × 10^-6^, and drug, F(1,24) = 2.0, *P* = 2 × 10^-4^, two-way within-subjects ANOVA); however, it had no effect in mice with bilateral GluN2B-SST KD (main effect of pre-pulse intensity, F(2,24) = 6.3, P = 1.7 × 10^-4^, but non-significant drug effect, F(1,12) = 4.0, P = 0.93; **Fig. 6c**). Bilateral GluN2B-SST KD in Cg1/M2 did not relieve all ketamine-induced abnormalities, as hyperlocomotion persisted (**Fig. 6d**). These results suggest that the effects of ketamine on some, but not all, behaviors rely on perturbing dendritic inhibition.

**Fig. 6.**
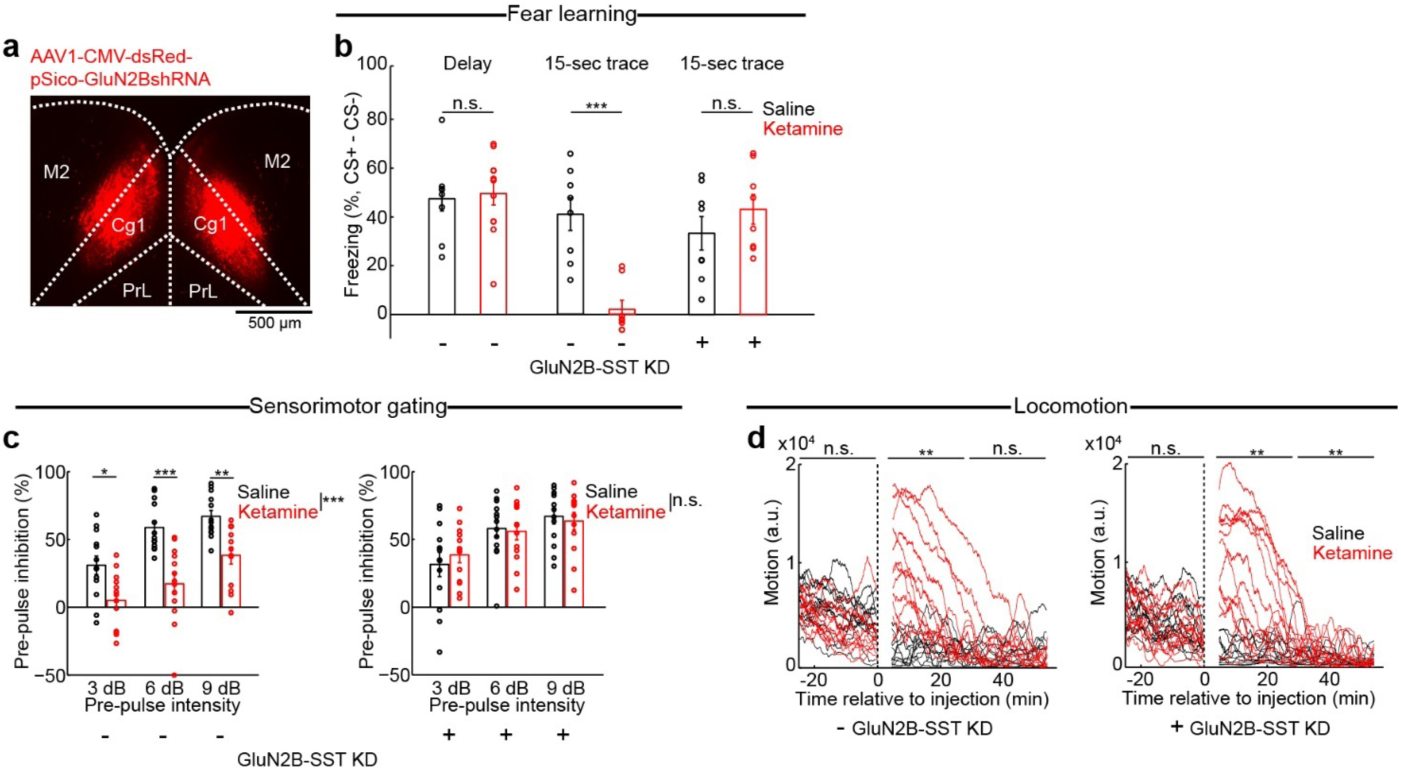
Chronic downregulation of GluN2B in prefrontal cortical SST interneurons and its behavioral consequences. (a) Coronal histological section, showing the extent of dsRed expression from bilateral injection of AAV1-CMV-dsRed-pSico-GluN2BshRNA into Cg1/M2 of a SST-IRES-Cre animal. (b) Fraction of time spent freezing during CS+ and trace period (if any), subtracted by that during CS- and trace period (if any), on the testing day. Freezing behavior for control SST-IRES-Cre animals in delay fear conditioning with no trace period (saline: 47 ± 6%, ketamine: 50 ± 5%, mean ± s.e.m.; *P* = 0.5, Wilcoxon rank-sum test; *n* = 8 and 12 animals for saline and ketamine respectively), in trace fear conditioning (saline: 41 ± 7%, ketamine: 2 ± 4%, mean ± s.e.m.; *P* = 6 × 10^-6^, Wilcoxon rank-sum test; *n* = 8 animals each for saline and ketamine), and for GluN2B-SST KD animals in trace fear conditioning (saline: 33 ± 7%, ketamine: 43 ± 6%, mean ± s.e.m.; *P* = 0.2, Wilcoxon rank-sum test; *n* = 8 animals each for saline and ketamine). (c) Pre-pulse inhibition as a measure of sensorimotor gating. For control SST-IRES-Cre animals, a two-way within-subjects ANOVA with pre-pulse intensity (3, 6, and 9 dB) and drug (saline, ketamine) as within-subjects factors found significant main effects of pre-pulse intensity (F(2,12) = 27.5, *P* = 4 × 10^-6^) and drug (F(1,24) = 2.0, *P* = 2 × 10^-4^), but a non-significant interaction (F(2,24) 1.8, *P* = 0.18). Post-hoc Tukey-Kramer’s tests for saline versus ketamine were significant at 3 dB (40 ± 7%, 5 ± 6%, mean ± s.e.m., *P* = 0.01), at 6 dB (59 ± 5%, 17 ± 8%, mean ± s.e.m., *P* = 9 × 10^-4^), at 9 dB (67 ± 4%, 38 ± 6%, mean ± s.e.m., *P* = 0.003). *n* = 13 animals each for saline and ketamine. For GluN2B-SST KD animals, two-way within-subjects ANOVA revealed a significant main effect of pre-pulse intensity (F(2,24) = 6.3, *P* = 1.7 × 10^-4^), but non-significant drug effect (F(1,12) = 4.0, *P* = 0.93) or two-way interaction (F(2,24) = 1.4, *P* = 0.26). Saline versus ketamine yielded no difference at 3 dB (32 ± 9%, 39 ± 6%, mean ± s.e.m.), at 6 dB (58 ± 6%, 56 ± 6%, mean ± s.e.m.), and at 9 dB (67 ± 6%, 63 ± 6%, mean ± s.e.m.). *n* = 13 animals each for saline and ketamine. (d) Open-field locomotor activity. Each trace comes from a single animal. For control SST-IRES-Cre animals, a two-way ANOVA was performed with epoch (pre-injection, 5-30 min post-injection, and 30-60 min post-injection) and drug (saline, ketamine) as within-subjects factors. There were significant main effects of epoch (F(2,22)=21.0, *P* = 7 × 10^-5^), drug (F(1,11) = 7.1, *P* = 0.02), and interaction (F(2,22) = 28.5, *P* = 7 × 10^-5^). Post-hoc Tukey-Kramer tests for saline versus ketamine were significant for 5-30 min post-injection (*P* = 0.001), but non-significant for pre-injection (*P* = 0.07) and 30-60 min post-injection (*P* = 0.58). *n* = 12 animals each for saline and ketamine. For GluN2B-SST KD animals, two-way ANOVA revealed significant main effects of epoch (F(2,22)=14.1, *P* = 0.002), drug (F(1,11) = 15.6, *P* = 0.002), and interaction (F(2,22) = 13.6, *P* = 0.003). Post-hoc Tukey-Kramer tests for saline versus ketamine were significant for 5-30 min post-injection epoch (*P* = 0.003) and 30-60 min post-injection (*P* = 0.001), but non-significant for pre-injection (*P* = 1.0). *n* = 13 animals each for saline and ketamine. * *P* < 0.05; ** *P* < 0.01; *** *P* < 0.001; n.s., not significant. Error bars, ± s.e.m.

Prefrontal GluN2B-SST KD reduced the activity of SST interneurons (**Fig. 5c**) and increased the rate of spontaneous synaptic calcium transients in pyramidal neurons (**Fig. 5e**). Although the manipulation rendered ketamine ineffective against trace fear learning and pre-pulse inhibition (**Fig. 6b, c**), GluN2B-SST KD alone had no detectable effect on these behaviors. Why does prefrontal GluN2B-SST KD reproduce ketamine’s effects on neural dynamics but not the behavioral outcomes? We hypothesized that the discrepancy may arise due to compensatory mechanisms. More specifically, behavioral testing occurred more than 4 weeks after GluN2B-SST KD, and it is possible that other brain regions become involved to restore behavioral functions^29^. To test this hypothesis, we performed experiments in which neural activity was reduced acutely using hM4D(Gi), a designer receptor exclusively activated by designer drug^30^ (DREADD) (**Fig. 7a**). After Cre-dependent expression of hM4D(Gi) in Cg1/M2 of SST-IRES-Cre mice, we treated the animals with ligand, clozapine-N-oxide (CNO), or vehicle and repeated the same trace fear conditioning and pre-pulse inhibition experiments along with ketamine or saline treatment. Consistent with the hypothesis, acute reduction of SST interneuron activity via CNO + hM4D(Gi) impaired trace fear learning and pre-pulse inhibition in saline-treated animals, while ketamine did not cause any additional effects (**Fig. 7b, c**). We performed control experiments to show that the effect was not due to hM4D(Gi) or CNO injection. Thus, unlike the chronic manipulations, an acute downregulation of prefrontal SST interneuron activity fully occludes the behavioral effects of ketamine on trace fear learning and pre-pulse inhibition.

**Fig. 7.**
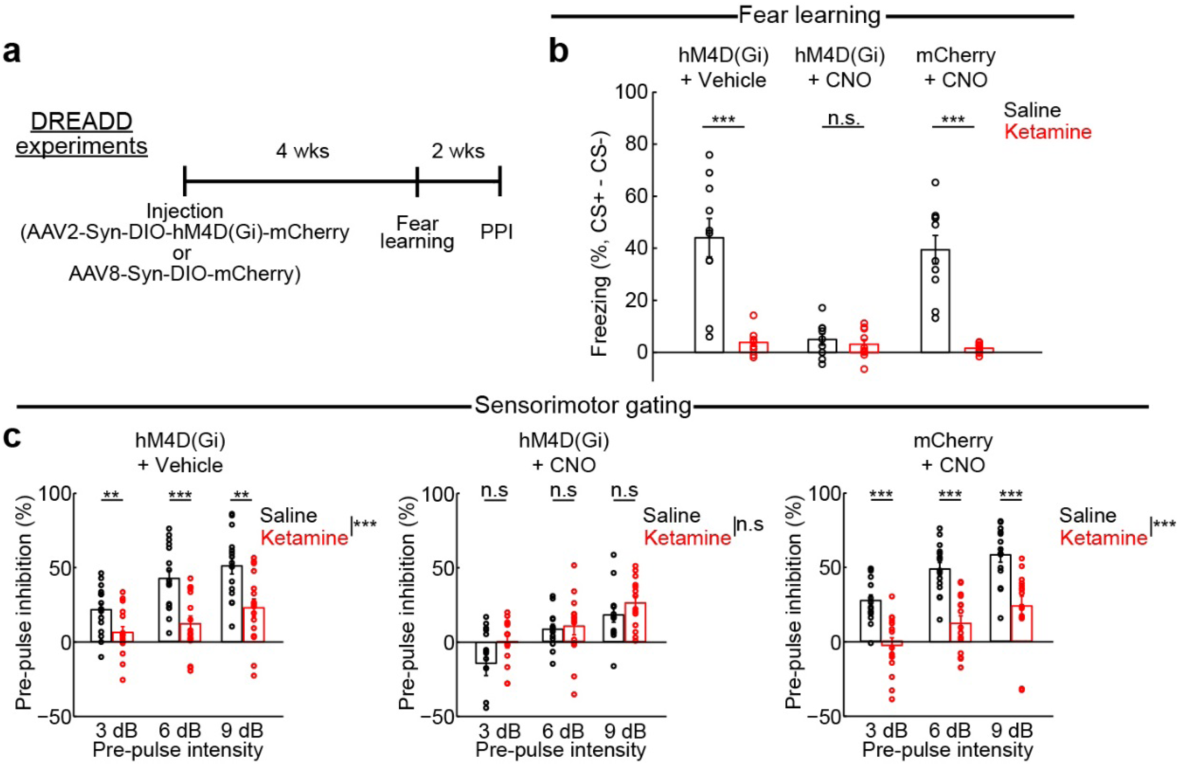
Acute suppression of SST interneuron activity in Cg1/M2 occludes the behavioral effects of ketamine. (a) Timeline for experiments involving use of Cre-dependent expression of designer receptors exclusively activated by designer drugs (DREADD) in SST-IRES-Cre animals. (b) Fraction of time spent freezing during CS+ and trace period, subtracted by that during CS- and trace period on the testing day. For hM4D(Gi) + Vehicle condition, ketamine treatment caused an impairment in freezing compared to saline (saline: 44 ± 7%, ketamine: 4 ± 2%, mean ± s.e.m.; *P* = 2 × 10^-4^, Wilcoxon rank-sum test; *n* = 10 animals for saline and 9 for ketamine). For hM4D(Gi) + CNO condition, there was impairment in freezing in saline-treated animals with no further difference relative to ketamine-treated animals (saline: 5 ± 2%, ketamine: 3 ± 2%, mean ± s.e.m.; *P* = 0.78, Wilcoxon rank-sum test; *n* = 9 animals for saline and 10 for ketamine). There was no effect of CNO on ketamine-induced impairment as tested in the mCherry + CNO condition (saline: 39 ± 5%, ketamine: 1.5 ± 1%, mean ± s.e.m.; *P* = 2 × 10^-4^, Wilcoxon rank-sum test; *n* = 10 animals each for saline and ketamine). (c) Pre-pulse inhibition as a measure of sensorimotor gating. For hM4D(Gi) + Vehicle condition, a two-way within-subjects ANOVA with pre-pulse intensity (3, 6, and 9 dB) and drug (saline, ketamine) as within-subjects factors found significant main effects of pre-pulse intensity (F(2,30) = 30.6, *P* = 6 × 10^-7^) and drug (F(1,15) = 21.9, *P* = 3 × 10^-4^), and a significant interaction (F(2,30) = 4.0, *P* = 0.03). Post-hoc Tukey-Kramer’s tests for saline versus ketamine were significant at 3 dB (22 ± 4%, 6 ± 4%, mean ± s.e.m., *P* = 0.01), at 6 dB (42 ± 6%, 12 ± 5%, mean ± s.e.m., *P* = 1 × 10^-4^), at 9 dB (51 ± 5%, 23 ± 6%, mean ± s.e.m., *P* = 0.002). *n* = 16 animals each for saline and ketamine. For hM4D(Gi) + CNO condition, a two-way within-subjects ANOVA found a significant main effect of pre-pulse intensity (F(2,26) = 11.9, *P* = 2 × 10^-4^) but a non-significant main effect of drug (F(1,13) = 4.1 *P* = 0.06) and interaction (F(2,26) = 1.2, *P* = 0.3). Post-hoc Tukey-Kramer’s tests for saline versus ketamine were not significant at 3 dB (−14 ± 8%, 1 ± 4%, mean ± s.e.m., *P* = 0.06), at 6 dB (8 ± 3%, 10 ± 6%, mean ± s.e.m., *P* = 0.71), at 9 dB (17 ± 5%, 26 ± 4%, mean ± s.e.m., *P* = 0.16). *n* = 14 animals each for saline and ketamine. For mCherry + CNO condition, a two-way within-subjects ANOVA found significant main effects of pre-pulse intensity (F(2,28) = 30.4, *P* = 9 × 10^-8^) and drug (F(1,14) = 56.0, *P* = 3 × 10^-5^), and a non-significant interaction (F(2,28) = 0.5, *P* = 0.62. Post-hoc Tukey-Kramer’s tests for saline versus ketamine were significant at 3 dB (27 ± 4%, -3 ± 5%, mean ± s.e.m., *P* = 5 × 10^-4^), at 6 dB (49 ± 4%, 12 ± 5%, mean ± s.e.m., *P* = 1 × 10^-5^), at 9 dB (58 ± 5%, 24 ± 7%, mean ± s.e.m., *P* = 1 ×10^-5^). *n* = 16 animals each for saline and ketamine.

### Ketamine-induced elevation of cortico-cortical connectivity depends on dendritic inhibition

Because we showed elevated synaptic calcium responses to long-range cortico-cortical inputs, an important question is whether the dendritic disinhibition reported here may contribute to abnormal functional connectivity. To measure communication between cortical regions, we recorded local field potentials (LFP) from Cg1/M2 and RSC (**Fig. 8a**). We characterized the ultra-slow fluctuations in the envelope of the gamma-band activity (**Fig. 8a, b**), which has been shown to correlate with blood-oxygenation-level dependent signals^31^. Reducing dendritic inhibition in Cg1/M2 led to elevated cortico-cortical coupling between Cg1/M2 and RSC, as evidenced by higher interareal correlation and coherence in GluN2B-SST KD compared to control SST-IRES-Cre animals (**Fig. 8c**). Ketamine, which normally also increased cortico-cortical coupling in control animals, had no further effect in mice with GluN2B-SST KD in Cg1/M2 (**Fig. 8d**). These data demonstrate that the deficient inhibitory control of dendritic compartments contributes to altered connectivity at the network level.

**Fig. 8.**
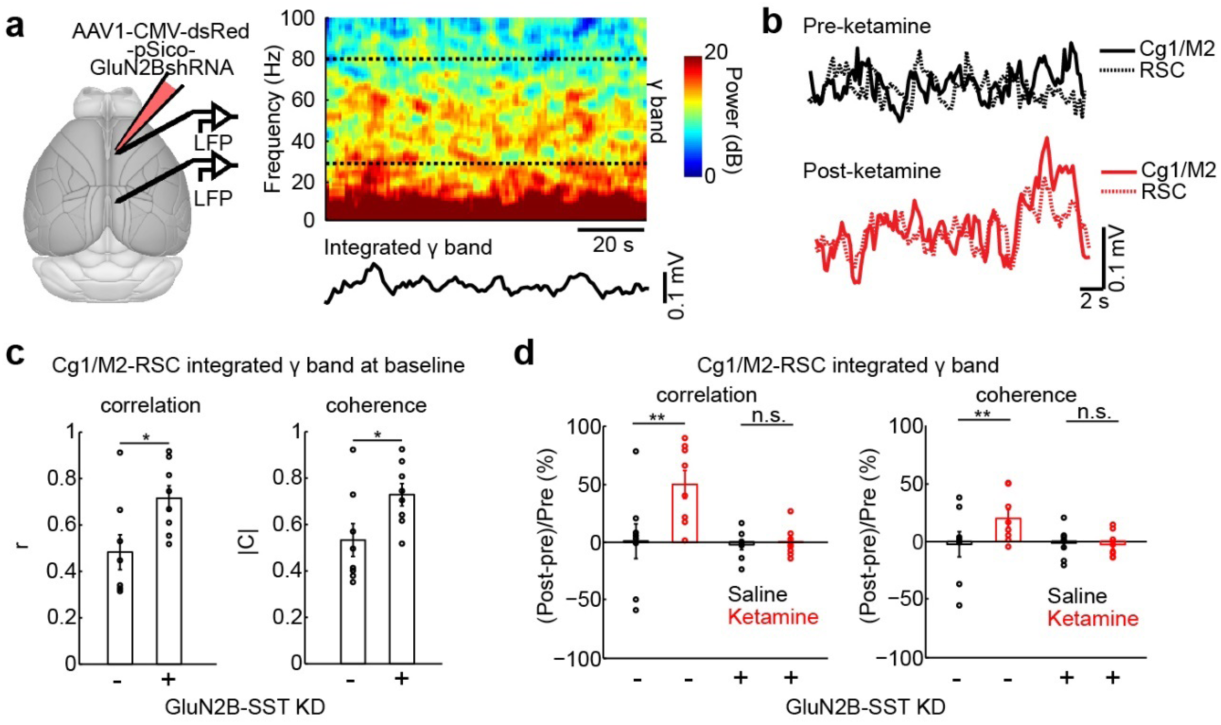
Elevated prefrontal cortical functional connectivity depends on dendritic inhibition. (a) Left, schematic of experimental setup. Right, spectrogram of an example local field potential (LFP) recording in Cg1/M2 of a SST-IRES-Cre animal. Dashed lines indicate the upper and lower bounds of the frequency range used to calculate the integrated gamma band signal, which is plotted below the spectrogram. (b) Integrated gamma band signals from LFPs recorded simultaneously from Cg1/M2 (solid line) and RSC (dashed line) of a SST-IRES-Cre animal. Example signals were plotted before (black) and after (red) ketamine injection. (c) Network-scale effects of prefrontal cortical SST interneuron manipulations (SST-IRES-Cre animals with or without GluN2B-SST KD). Functional connectivity between Cg1/M2 and RSC was quantified by determining the correlation and coherence between integrated gamma band signals of the two brain regions. At baseline prior to injection, control and GluN2B-SST KD animals had significantly different correlation coefficients (controls: 0.48 ± 0.08, GluN2B-SST KD: 0.72 ± 0.05, mean ± s.e.m.; *P* = 0.03, Wilcoxon rank sum test) and coherence magnitude (controls: 0.53 ± 0.07, GluN2B-SST KD: 0.73 ± 0.05, mean ± s.e.m.; *P* = 0.04, Wilcoxon rank sum test). *n* = 8 animals each for controls and GluN2B-SST KD. (d) Functional connectivity between Cg1/M2 and RSC after ketamine or saline for SST-IRES-Cre animals with or without GluN2B-SST KD. For correlation, a two-way mixed ANOVA was performed with treatment (GluN2B-SST KD, no KD) as the between factor, and drug (saline, ketamine) as the within factor. The interaction effect was significant (F(1,14) = 6.2, *P* = 0.03). Post-hoc Tukey-Kramer test showed a significant ketamine-saline difference in the no KD group (*P* = 0.003), but not in the GluN2B-SST KD group (*P* = 0.9). For coherence, the interaction effect was significant (F(1,14) = 7.1, *P* = 0.02). Post-hoc Tukey-Kramer test showed a significant ketamine-saline difference in the no KD group (*P* = 0.003), but not in the GluN2B-SST KD group (*P* = 0.8). *n* = 8 animals each for controls and GluN2B-SST KD. * *P* < 0.05; ** *P* < 0.01; *** *P* < 0.001; n.s., not significant.

## Discussion

The goal of this study was to delineate the impact of subanesthetic ketamine on the prefrontal cortical circuit. There were two major findings. One, ketamine reduces the activity of dendrite-targeting SST interneurons. Two, as a consequence of the diminished dendritic inhibition, ketamine increases the amplitude of the synaptically evoked calcium transients in the apical dendritic spines of pyramidal neurons. As we will elaborate below, these local-circuit alterations provide insights into the neural mechanisms for the acute psychotomimetic and sustained antidepressant effects of ketamine.

The novelty of our results is to highlight SST interneurons as a key target of systemic NMDAR antagonism. Although in principle the disinhibition hypothesis is general and need not be tied to particular GABAergic neuron subtypes, so far empirical studies of NMDAR hypofunction have focused on fast-spiking cells for technical reasons. For example, recordings of inhibitory postsynaptic currents have provided clear evidence for disinhibition^32, 33^, but the method cannot distinguish the cell types that provide the GABAergic inputs, and somatic electrodes tend to favor the detection of perisomatic inputs. *In vivo* electrophysiology has also supported disinhibition by showing reduced activity for fast-spiking cells^12^, but again SST interneurons are likely not examined in this earlier study because their spike waveforms have overlapping features with excitatory neurons and thus cannot be easily isolated^17^. By contrast, using a GABAergic Cre-driver mouse line, our fluorescence signals would originate predominantly from SST interneurons. The ability to characterize responses from genetically identified SST interneurons distinguishes our results from prior studies of ketamine and neural activity.

Perisomatic and dendritic disinhibition may represent separate, parallel ways by which ketamine acts on the prefrontal cortex. This is probable because when we manipulated dendritic inhibition, the behavioral and electrophysiological alterations are largely orthogonal to those previously reported for NMDAR manipulations in fast-spiking cells. More specifically, when the obligatory GluN1 subunit was deleted from fast-spiking cells, behavioral deficits were reported for delay fear conditioning, but not for pre-pulse inhibition^34^. This is opposite to what we saw. In another study with postnatal GluN1 removal in many fast-spiking cells, although here there was a deficit in pre-pulse inhibition similar to our animals, the NMDAR antagonist-induced locomotion was reduced^35^, unlike our manipulation of SST interneurons which had no effect on the ketamine-induced hyperlocomotion. Finally, enhanced baseline gamma-band oscillations were observed following NMDAR deletion in fast-spiking cells^34, 36^ and may even be produced by removing NMDARs in pyramidal neurons^37^, whereas we show that dendritic inhibition was involved in controlling the envelope of oscillatory signals on a much slower timescale. This interpretation comes with the caveat that we are comparing behavioral outcomes from studies involving NMDAR manipulations to different extent and at different developmental time points. Notwithstanding the caveat, it seems possible that ketamine’s suppressive effects on perisomatic and dendritic inhibition may be responsible for non-overlapping sets of behavioral and electrophysiological phenotypes.

We show that the ketamine-induced loss of dendritic inhibition leads to heightened synaptic calcium transients in the dendritic spines of pyramidal neurons. This is consistent with what we know about SST interneurons: they regulate dendritic excitability and control the amount of calcium influx in dendritic spines in prefrontal cortical brain slices^38, 39^. In an awake mouse, the inhibitory influence of SST interneurons is expected to be greater, as the cells have high spontaneous firing rates *in vivo*^40^, setting the stage for the suppression by ketamine. The positive skew in the ketamine-induced increase of synaptic calcium transients (**Fig. 2d**) suggests that only a subset of dendritic spines was impacted by ketamine. This agrees with anatomical studies, showing that only ∼20 – 50% of cortical dendritic spines are innervated by SST axonal boutons^38, 41, 42^. By contrast, for SST interneuron soma (**Fig. 1i**) and axonal boutons (**Fig. 2b**), ketamine appears to induce a subtractive shift of the entire activity distribution, which is consistent with most SST neurons having NMDAR-mediated currents^26^. One intriguing observation is that the synaptic calcium elevations were observed in the medial prefrontal cortex, but not the primary motor cortex. This may be because although the systemic administration is expected to deliver ketamine to all brain regions, the density of cortical SST interneurons is highest in the frontal associations regions^43^. The gradient of cortical interneuron density may explain the heterogeneous, region-specific responses to a global perturbation, as has been proposed recently^44^.

How might the dendritic disinhibition relate to the behavioral effects of subanesthetic ketamine? Acutely, ketamine produces psychotomimetic symptoms, perceptual aberrations, and cognitive impairments. In the mouse, the Cg1/M2 sub-regions of the medial prefrontal cortex are involved in linking antecedent events to current decisions^45^. Dendritic inhibition is thought to be crucial for the context-dependent gating of inputs^46^. Dysregulation of dendritic inhibition is therefore expected to allow aberrant sensory inputs into the prefrontal cortex, thereby disorganize the task-specific activity patterns of prefrontal cortical neurons^47^. Experimentally, this matches with our finding of ketamine-induced prefrontal cortical hyperconnectivity. Our mouse experiments were motivated by human neuroimaging studies that have found abnormally high functional connectivity for the prefrontal cortex in humans with ketamine or early-course schizophrenia^48, 49^. Crucially, here we have extended to test the potential neural mechanism by showing that the prefrontal cortical hyperconnectivity in the mouse depends on the ability of ketamine to perturb dendritic inhibition.

What about the antidepressant effects of subanesthetic ketamine? One striking consequence of ketamine is the synaptic adaptations in the medial prefrontal cortex that can be detected 24 hr later, including increased dendritic spine density and strengthened synaptic responses^50, 51, 52^. It is thought that these positive influences on synapse function might counteract the impaired synaptic transmission observed in rodents after chronic stress^53^ and the reduced synaptic connections in stressed rodents and depressed subjects^54, 55^. One possibility is that the heightened calcium transients in dendritic spines within the first hour of systemic administration is important for synaptic plasticity. A potential link between the acute calcium elevations and subsequent synaptic strengthening is supported by studies that have shown postsynaptic calcium influx to be required and sufficient to induce activity-dependent synaptic plasticity^56, 57^. It agrees with the observation that the antidepressant effect is blunted if ketamine is administered while excitatory transmission is blocked^50, 58^, and that manipulations of SST interneurons can tune depressive-like behaviors in rodents^22, 59^. SST interneurons are also known to connect densely with pyramidal networks^60^ and thereby regulate their correlated firing^61^, which may relate to a very recent observation that ketamine restores coordinated activity in prefrontal ensembles^52^. Therefore, there are reasons to suspect that dendritic disinhibition might be relevant for ketamine’s antidepressant effects. Additional studies will be required to more firmly establish the potential relationship.

## Methods

### Animals

Adult male mice (>10 weeks of age at the start of experiments) of the C57BL/6J strain (Stock No. 000664, Jackson Laboratory) were used. To selectively image or manipulate SST interneurons, we utilized adult male SST-IRES-Cre^62^ on a mixed C57BL/6 × 129S4/SvJae background (Stock No. 013044, Jackson Laboratory). Mice were group housed (3-5 mice per cage) on a 12:12-hr light/dark cycle with free access to food and water. We typically performed experiments during the light cycle. Sample sizes for this study are selected based on numbers generally employed in the field. All experimental procedures were approved by the Institutional Animal Care and Use Committee at Yale University.

### Surgery

For all stereotaxic surgical procedures, the mouse was first anesthetized with isoflurane in oxygen (3-4% during induction, 1-1.5% for the remainder of the surgery) while placed in a stereotaxic apparatus (David Kopf Instruments). The animal lay on top of a water-circulating pad (Gaymar Stryker) set at a constant temperature of 38 °C. Eyes were lubricated with ophthalmic ointment. Carprofen (5 mg/kg, s.c.; 024751, Henry Schein Animal Health) and dexamethasone (3 mg/kg, i.m.; 002459, Henry Schein Animal Health) was used pre-operatively. The skin above the skull was then sterilized by swabs of ethanol and betadine before an incision was made to expose the skull. We made a craniotomy (small or large depending on the experiment as detailed below) using a dental drill. Immediately after surgery and on each day for the following 3 days, carprofen (5 mg/kg, s.c.) was given to the animal. All animals had at least 1 week to recover before starting experiments unless stated otherwise.

#### For imaging

We made a small skin incision and craniotomy (∼0.5 mm diameter) in the right hemisphere centered at 1.5 mm anterior of bregma (anterior-posterior, AP) and 0.3 mm lateral of midline (medial-lateral, ML) to target the medial prefrontal cortex (Cg1 and M2 sub-regions)^63^. A glass pipette (3-000-203-G/X, Drummond Scientific) was pulled to a fine tip (P-97 Flaming/Brown Micropipette Puller, Sutter Instrument), front-filled with the relevant viruses (see below) and then slowly lowered into the brain using the stereotaxic apparatus at 4 sites corresponding to vertices of a 0.1-mm wide square centered at the above coordinates. At depth of 0.3 to 0.5 mm below the dura, we ejected the virus using a microinjector (Nanoject II, Drummond Scientific). To target the retrosplenial cortex (RSC) and primary motor cortex (M1), we used the same approach centering at AP = -1.4 mm/ML = 0.3 mm, and AP = 1.8 mm/ML = 1.6 mm respectively. We injected 9.2 or 18.2 nL per injection pulse, with ∼30 s in between each injection pulse. To reduce backflow of the virus, we waited at least 5 minutes after finishing injection at one site before retracting the pipette to move on to the next site. For experiments that required more than one virus, we injected each virus sequentially. The brain was kept moist with artificial cerebrospinal fluid (ACSF, in mM: 5 KCl, 5 HEPES, 135 NaCl, 1 MgCl2, 1.8 CaCl2; pH 7.3). After completing all injections, the craniotomy was covered with silicone elastomer (0318, Smooth-On, Inc.), and the skin sutured (Henry Schein). After approximately 3 weeks, the animal underwent a second surgery for cranial window implant. After the initial steps of anesthesia as outlined above, an incision was made to remove all of the skin above the skull, which was cleaned to remove connective tissues. We made a 3-mm diameter craniotomy around the previously targeted location. A glass window was put together by gluing two 3-mm-diameter, #1-thickness circular glass (640720, Warner Instruments), using UV-sensitive optical adhesive (NOA61, Norland Products). The glass window was then placed on the exposed brain with slight downward pressure while applying high-viscosity adhesive (Loctite 454) around the perimeter of the glass window. After the adhesive cured, a custom-made stainless steel headplate (eMachineShop.com) was affixed to the skull using quick adhesive cement (C&B Metabond, Parkell).

#### For behavior

We followed the surgical procedures for imaging, with the following exceptions. There was no second surgery for cranial window and head plate implant. All injections were bilateral injections, in which we used one of the coordinates above, plus the mirrored coordinates for the other hemisphere. We injected at all the locations in one hemisphere first, before front-filling more virus (if necessary), and moving onto the other hemisphere. We waited at least 4 weeks before behavioral testing.

#### For local field potential recordings

We fabricated bundles of stainless steel wire electrodes (2-3 wires per bundle) (790500, A-M Systems). The diameter of an electrode was 114.3 µm and 50.8 µm with and without coating respectively. There was ∼350 µm of exposed tip for each electrode which was separated by ∼250 µm from other electrodes in the bundle. Impedance was typically less than 100 kΩ when tested at 1 kHz. Following the procedures above for surgery and small craniotomy, a bundle of electrodes was each lowered into the right Cg1/M2 and right RSC, centered at the stereotaxic coordinates as described above, to a depth of 0.5 mm below the dura. We used a miniature stainless steel screw (#0000-160) placed over the right cerebellum as the reference electrode. All electrodes were then soldered to a miniature connector (A79002-001 or A79000-001, Omnetics). Care was taken to avoid shorted connections. All pieces of the implant along with a stainless steel headplate were affixed to the skull using cyanoacrylate (Loctite 454) and quick adhesive cement (C&B Metabond, Parkell). For experiments combining viral injections and electrophysiology, we first injected the virus, and then waited 3 weeks before proceeding with a second surgery for electrode implant.

#### For electrical stimulation

We first injected viruses using the procedures above. Three weeks later, we performed a second surgery to implant stimulating electrodes. A bundle of stainless steel wires (3-4 wires per bundle) (790500, A-M Systems) was inserted into RSC. The wire implant was soldered to a miniature connector (A79002-001 or A79000-001, Omnetics). A glass-covered cranial window was made over Cg1/M2 using procedures described above. All pieces of the microstimulation implant and a stainless steel headplate were affixed to the skull using cyanoacrylate (Loctite 454) and quick adhesive cement (C&B Metabond, Parkell).

### Histology

Mice were transcardially perfused with paraformaldehyde solution (4% (v/v) in phosphate-buffered saline). The brains stayed in the fixative for at least 48 hours, and then were sectioned coronally with a vibratome and mounted on slides with coverslips. We imaged the sections with an upright fluorescence microscope (Zeiss Axio Imager M2).

### Viruses

To image calcium transients in pyramidal neurons and their dendrites, we injected AAV1-CamKII-GCaMP6f-WPRE-SV40 (Penn Vector Core) diluted to a titer of 2 × 10^12^ genome copies (GC) per milliliter. A total of 73.6 nL of virus was injected for each experiment. For interneurons, we injected AAV1-Syn-DIO-GCaMP6s-WPRE-SV40 (Penn Vector Core; 1 × 10^12^ GC/mL titer, 147.2 nL) in SST-IRES-Cre animals. To downregulate GluN2B expression in a cell-type-specific manner, we designed a shRNA sequence targeting mouse GluN2B (5’ TGTACCAACAGGTCTCACCTTAAACTTCAAGAGAGTTTAAGGTGAGACCTGTTGGTACTTTTTTC 3’), ligated the GluN2BshRNA sequence into a modified pSico plasmid backbone as previously described ^22^, and packaged the plasmid into an AAV to create AAV1-CMV-dsRed-pSico-GFP-GluN2BshRNA. For behavioral experiments, a total of 966 nL was injected in Cg1/M2 of each hemisphere as well as at additional sites around Cg1/M2 in the anterior-posterior axis. For imaging experiments, the GluN2BshRNA sequence was ligated into another pSico backbone with no GFP (Vector Biolabs; AAV1-CMV-dsRed-pSico-GluN2BshRNA, 2 × 10^13^ GC/mL titer). The reason for removing the GFP is to avoid interference with GCaMP6f fluorescence. A total of 552 nL was injected in Cg1/M2, and then followed by injection of AAV1-CamKII-GCaMP6f-WPRE-SV40. For electrophysiology experiments, a total of 552 nL of AAV1-CMV-dsRed-pSico-GluN2BshRNA was injected in Cg1/M2. To acutely downregulate SST interneuron activity, we injected either AAV2-Syn-DIO-hM4D(Gi)-mCherry (Penn Vector Core, 1 × 10^13^ GC/mL titer, 552 nL) for the hM4D(Gi) group or AAV8-Syn-DIO-mCherry (Penn Vector Core, 1 × 10^13^ GC/mL titer, 552 nL) for the control group in SST-IRES-Cre animals.

### Cell culture validation

We used cell cultures to validate the viral-mediated manipulation of NMDAR expression. Primary neuronal cultures were prepared as previously described ^64^. Briefly, cortical neurons at E18 from pregnant female rats were dissected, dissociated and plated in wells (∼1 million cells/well) and cultured in standard media. We added viruses at DIV5 stage. For GluN2BshRNA validation, we used the viruses AAV1-CMV-Cre-GFP (1 × 10^13^ GC/ml titer, 400 nL) for expression of Cre and AAV1-CMV-dsRed-pSico-GluN2BshRNA (2 × 10^13^ GC/ml titer, 400 nL) for expression of GluN2BshRNA. Using these two viruses, we made four well conditions: GluN2BshRNA, no Cre; GluN2BshRNA, Cre; no GluN2BshRNA, Cre; no GluN2BshRNA, no Cre (no virus added) (**Supplementary Fig. 5**). We then performed Western blots with antibodies for GluN2B (#4207S, Cell Signaling, 1:1000, rabbit polyclonal antibody) and beta-actin (#4967S, Cell Signaling, 1:1000, rabbit polyclonal antibody). Bands were quantified using Bio-Rad Image Lab software and raw band intensity values were normalized to beta-actin levels.

### Drugs

We used saline (0.9% Sodium chloride, Hospira). Ketamine working solution (1 mg/mL in saline) was prepared from stock solution (McKesson), approximately every 30 days. Clozapine N-oxide (Sigma Millipore) working solution (1 mg/ml) was prepared fresh daily before each experiment from a stock solution dissolved in DMSO (10 mg/ml) stored at 4 degrees C. Vehicle (1:10 DMSO in saline) was used in control injections.

### Two-photon calcium imaging

We used a laser-scanning two-photon microscope (Movable Objective Microscope, Sutter Instrument) controlled by the ScanImage software^65^. A tunable Ti:Sapphire femotosecond laser (Chameleon Ultra II, Coherent) was the excitation source which was focused onto the brain using a water immersion objective (XLUMPLFLN, 20X/0.95 N.A., Olympus). The time-averaged laser power was typically less than 100 mW during experiments. Filters had the following center excitation (ex) and emission (em) wavelengths (Semrock or Chroma): GCaMP6s/f, 920 nm (ex) and 525 nm (em); dsRed, 1040 nm (ex) and 605 nm (em). Emitted photons were collected by GaAsP photomultiplier tubes. We acquired images that are 256 × 256 pixels at 0.68 or 0.54 µm per pixel resolution for dendritic spines and axonal boutons or 1.35 µm per pixel resolution for cell bodies. Frame rate was 3.62 Hz using bidirectional scanning on a set of galvanometer-based scanners. A software program (Presentation, NeuroBehavioral Systems) sent TTL pulse via a data acquisition device (USB-201, Measurement Computing) to the two-photon microscope at 30-s intervals, as well as to trigger all the other equipment. The Presentation software would log timestamps so that data from imaging and other equipment (e.g., microstimulation and body motion) can be synchronized in later analyses. Mice were habituated to head fixation under the microscope for 3 - 5 days of increasing durations before the commencement of any data collection. While head fixed, the mouse would sit in an acrylic tube, which permits postural adjustments but limits gross body movements. Subcutaneous injections were given through a drilled hole in the tube so as to allow the mouse to remain head fixed. To examine the acute effects of ketamine, we would image a field of view for 10 – 15 minutes to obtain ‘pre-injection’ data. The imaging was then interrupted for ketamine (10 mg/kg) or saline (10 mL/kg) injection (s.c.). At 30 minutes after the injection, we would verify that the same field of view was still in focus, and then obtain imaging data for another 10 – 15 minutes. Imaging 30 minutes after the injection ensured that we bypass the transient hyperlocomotion effects of ketamine (**Fig. 1c, d**). In a subset of animals, we imaged continuously for 60 min after injection, but analyzed only the data after the 30-minute post-injection mark. Each subject was imaged for ketamine and saline injections, with the order counterbalanced across subjects. We imaged at 0 – 200 µm below dura for dendritic spines, and 200 – 500 µm below dura for cell bodies. We targeted the medial portion of Cg1/M2, within 0 – 500 µm of the midline as visualized by the dark band of the midsagittal sinus. For imaging experiments with cell-type-specific knockdown experiments, during live imaging, we searched for and imaged fields of view with many co-labeled cells that expressed dsRed.

### Analysis of calcium imaging data

We first concatenated all of the .tiff image files from the same experiment and corrected for lateral motion using either TurboReg implemented as a plugin in ImageJ or NoRMCorre^66^ in MATLAB. A custom graphical user interface (GUI) in MATLAB was used for the manual selection of region of interests (ROI). The rest of the analysis depended on experiment type as described below.

#### Dendritic spines

We manually scrolled through the imaging frames to look for ROIs that likely represented dendritic spines and dendritic shaft – neurite segments with multiple protrusions that display correlated patterns of fluorescence transients. The dendritic shaft ROIs were selected to be within 20 µm of the corresponding dendritic spine ROIs. We averaged values of pixels within a ROI to generate *F*ROI(*t*). For each ROI, we quantified the contribution of neuropil to the fluorescence signal. Specifically, we would calculate a radius, *r*, treating the ROI area as the area of a circle, and then creating an annulus-shaped neuropil ROI with inner and outer radii of 2*r* and 3*r* respectively. We excluded pixels that belonged to the ROIs of other spines and shaft. To also exclude pixels that may belong to unselected dendritic structures, we calculated the time-averaged signal for each pixel, and then determined the median for all pixels within the neuropil ROI. We then excluded pixels if their time-averaged signal was higher than the median. We finally averaged across the non-excluded pixels in the neuropil ROI to generate *F*_neuropil_(*t*). To subtract the neuropil signal, we used the formula:

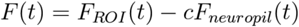

where *c* = 0.4 is the neuropil correction factor. The fractional change in fluorescence, Δ*F*/*F*(*t*), for each spine and shaft was calculated as follows:

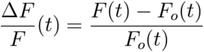

where *F*_0_ (*t*) is the 10^th^ percentile of *F*(*t*) within a 2-minute sliding window. Because we were presently interested in calcium signals due to subthreshold synaptic inputs, we regressed out fluorescence signals from non-local sources based on a previously described procedure^18^. Briefly, for each spine with Δ*F*/*F*_spine_(*t*), we computed Δ*F*/*F*_synaptic_(*t*) by subtracting out the scaled version of fluorescence transients in the corresponding dendritic shaft, Δ*F*/*F*_shaft_(*t*), using the following equation:

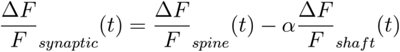

where α was determined by a linear regression forced through the origin of Δ*F*/*F*_spine_(*t*) versus Δ*F*/*F*_shaft_(*t*).

#### Axonal boutons

We drew ROIs that are likely single axonal boutons – varicosities on a relatively thin neurite branch with no protrusions. This manual identification is supported by the known neuroanatomy. For SST interneurons, they send axons but very few dendrites, if any, to layer 1^67^. For retrosplenial cortical neurons, they send long-range axons to layer 1 of Cg1/M2. To generate *F*_ROI_(*t*), values of pixels within an ROI were averaged. For each ROI, we would estimate the contribution of neuropil to the fluorescence signal, by calculating and subtracting *F*_neuropil_(*t*) to obtain *F*_candidate_bouton_(*t*), using the procedures described in the last section. Some boutons may belong to the same neuron. To limit duplicates, we used a modified version of a previously published procedure^68^. Briefly, we first analyzed a subset of imaging experiments to identify pairs of boutons on the same branch, based on distance (within 10 µm of each other) and visual confirmation of a connecting axonal segment. From this subset of bouton pairs (*n* = 46 bouton pairs, 2 animals), we used their *F*_candidate_bouton_(*t*) values to compute a correlation coefficient for each pair. The 5^th^ percentile value for all the correlation coefficients in this subset was 0.59, which was used as the cut-off threshold value. Then, for each axonal imaging experiment, we performed an iterative procedure to select unique boutons. In each iteration, a correlation matrix using *F*_candidate_bouton_(*t*) between all pairs of candidate boutons in a field of view was computed. The pair of candidate boutons with the highest correlation coefficient was identified. If the value exceeded the threshold of 0.59, we would select one out of that pair of candidate boutons randomly, and looked for other boutons whose correlation coefficient with the selected bouton exceeded the threshold of 0.59. The original identified pair and boutons from subsequent search then would form one cluster. Entries in the correlation matrix corresponding to this cluster would be excluded for subsequent iterations. The process continued again to identify another cluster until no pair of candidate boutons in the correlation matrix had value above the threshold of 0.59. At the end of the iterative procedure, each of the remaining boutons was assigned to be its own cluster. We randomly selected one bouton as the representative for each cluster. The fluorescence signals of these representative boutons, *F*_bouton_*(t)*, were then converted to fractional changes in fluorescence, Δ*F*/*F*_bouton_(*t*), as described above and used for subsequent analyses.

#### Cell bodies

We selected ROIs that are likely cell bodies – fluorescent objects with a round boundary and a lateral extent of about 10 µm. Values of pixels within a ROI were averaged to generate *F*_ROI_(*t*). For each ROI, we would remove the contribution of neuropil to the fluorescence signal, by calculating and subtracting *F*_neuropil_(*t*) to obtain *F*_soma_(*t*), and then calculating the fractional changes in fluorescence Δ*F*/*F*_soma_(*t*), using procedures outlined in the previous sections. For experiments with cell-type-specific GluN2B manipulation, we selected only GCaMP6s-expressing cells that were co-labeled with dsRed.

#### Detection of calcium events

After obtaining Δ*F*/*F*_synaptic_(*t*), Δ*F*/*F*_bouton_(*t*), or Δ*F*/*F*_soma_(*t*) for a ROI, we did the following analysis to detect calcium events. We employed a previously characterized and validated “peeling” algorithm^16, 69^, which used an iterative, template-matching procedure to decompose Δ*F*/*F*(*t*) into a series of elementary calcium events. Briefly, we used a template for an elementary calcium event instantaneous onset with an amplitude of 0.3, and a decay time constant of 1 s for a single-exponential function. The algorithm would search for a match to this specified template in Δ*F*/*F*(*t*). When a match occurred, a calcium event was recorded, and the template subtracted from Δ*F*/*F*(*t*) (i.e., peeling). The remaining Δ*F*/*F* trace was then searched for another match, in an iterative manner until no further matches were found. The times of recorded calcium events formed the output of the algorithm. The temporal resolution of the output is limited by the imaging frame rate. There can be multiple events associated with the same event time. In the original paper describing the peeling algorithm, extensive numerical simulations suggested excellent detection performance, even in worst-case scenarios with low signal-to-noise ratios and low sampling rate^16^. For each imaging session, we calculated calcium event rate by dividing the number of calcium events by the duration of the imaging period. For **Fig. 2e**, we characterized the calcium events in terms of amplitude and frequency. We identified calcium events that co-occurred in the same imaging frame (same event time), distinguishing amplitude (mean number of calcium events per frame, for frames with at least one event) from frequency (number of frames with at least one event divided by the duration of the imaging session). We imaged the same compartment (spine, bouton, or cell body) before and after injection, and therefore were able to quantify the change in calcium event rate, amplitude and frequency for each compartment by calculating the post-minus pre-injection values normalized by the pre-injection value.

#### Detection of calcium events using an alternative method

To determine if our results depend on the event detection procedure, we used an alternative method to quantify calcium transients in dendritic spines (**Supplementary Fig. 3**). We started from Δ*F*/*F*_synaptic_(*t*). For each ROI and for each condition (i.e., pre- or post-injection), we would calculate a threshold corresponding to 3 times the median absolute deviation of Δ*F*/*F*_synaptic_(*t*). We noted imaging frames in which Δ*F*/*F*_synaptic_(*t*) was above the threshold. From this subset of frames, we determined the mean amplitude above threshold (Δ*F*/*F*_synaptic_(*t*) minus the threshold), and the fraction of time spent above the threshold (the number of above-threshold frames divided by the number of all frames) (see **Supplementary Fig. 3a** for an illustration). We quantified the changes in these measures - mean amplitude above threshold and fraction of time above threshold - by calculating the post-minus pre-injection values normalized by the pre-injection value.

### Recording animal movements during two-photon imaging

To assess the mouse’s movements during two-photon imaging (**Fig. 1c, d**), we illuminated the animal with infrared light and recorded videos (640 x 480 pixels, 20Hz) of the animal’s face and body using an infrared camera (See3CAM_12CUNIR, e-con Systems). Although the animal was head-fixed, it could still move parts of its body such as the face, trunk, and limbs. To synchronize the infrared video with imaging, the infrared illuminator was triggered to turn off for 100 ms by the same TTL pulse used to trigger the two-photon microscope. During analysis, the interval between dark frames therefore indicated a duration of 30 s. We computed motion *m* in 1-s bins:

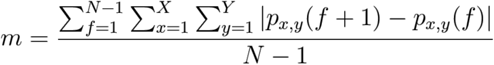

where *N* is the number of video frames in the 1-s bin, X and Y are the width and height of the frame, *p*_*x,y*_*(f+1)* and *p*_*x,y*_*(f)* are values of the pixel at coordinates *x* and *y* in frames *f+1* and *f* respectively.

### Simultaneous electrical stimulation and imaging

Imaging in Cg1/M2 was performed as described above. For microstimulation, we injected electrical currents into the implanted stimulation electrodes in RSC, using an analog stimulus isolator (Model 2200, A-M Systems). The timing and strength of the stimulation were specified by a script written in a software program (Presentation, NeuroBehavioral Systems) via command voltages out of a multifunctional input/output device (USB-6001, National Instruments). During each experiment, to calibrate the stimulation current (which could vary due to depleting battery for the stimulus isolator or high impedance of the implanted electrode), an electronic circuit was placed in-series, allowing us to measure the stimulation current online and tune the electrical current to within 5% of the desired level. For the stimulation pattern, each pulse had a biphasic waveform, lasted 5 ms in each phase, and had peak amplitudes of ±150 µA. We used 1, 2, 4, 8, 16, 32 or 64 pulses as the stimulation levels. If more than 1 pulse was used, the pulses were applied one after the other with no inter-pulse interval. A block consisted of the 7 stimulation levels, presented in random order with a random inter-trial period of 10 – 14 s. For each experiment, we presented 10 blocks successively before injection of ketamine or saline, and then presented another 10 blocks successively 30 – 60 minutes after injection. Prior to the post-injection stimulation, we imaged to confirm that the field of view has not changed. Using procedures described in earlier sections, we determined calcium events in each dendritic spine and axonal bouton. The log files generated by the Presentation software were used to align the timestamps of the imaging data to the stimulation events. For each dendritic spine or axonal bouton, we determined the number of calcium events detected within 1 s following the onset of stimulation, and reported the averaged values across the 10 blocks.

### Electrophysiology

Animals were head-fixed and placed in a Faraday cage. An interface cable was mated to the connector on the animal’s head in order to amplify local field potential (LFP) signals using a differential amplifier (RHD2216, Intan Technology), high-pass filtered at 0.5 Hz and digitized at 1 kHz (RHD2000, Intan Technology). Before actual electrophysiological recordings, mice were habituated to head-fixation for 3 – 5 days of increasing durations. We recorded for 30 minutes pre-injection, and then for 60 minutes post-injection. We discarded data from electrodes that were noisy (impedance >1 GΩ). One electrode from each site (Cg1/M2 or RSC) was chosen randomly for analysis. For analysis, we compared the 30-minute segment recorded pre-injection to the 30-minute segment recorded from 30 – 60 minutes post-injection. We focused on the integrated gamma band power, because it was previously found to correlate with hemodynamic signals underlying blood-oxygenation-level-dependent resting-state connectivity^31^. To determine the integrated gamma band power, spectrograms were computed from the local field potential recordings using a sliding window with duration of 2 s and step size of 0.4 s. At each window, spectral power within the gamma band from 30 to 80 Hz was summed, and this yielded the integrated gamma band power. Correlation between integrated gamma band power in Cg1/M2 and RSC was calculated based on Pearson correlation coefficient at zero lag. Coherence between integrated gamma band power in Cg1/M2 and RSC was computed using the Chronux^70^ package in MATLAB, and the magnitude of coherence was reported.

### Behavior

#### Fear learning

Fear learning was assessed using a conditioning box utilizing an infrared video camera (320 × 240 pixels, 30 Hz; Med Associates, Inc.). To enable more flexible protocols involving multiple stimuli and repetitions, we modified the box to be controlled by a behavioral control software (Presentation, NeuroBehavioral Systems). Each of the conditioned stimuli (CS) was a 20-second long series of auditory pips (500 ms on, 500 ms off). The pip was either 2.5 kHz (85 dB, calibrated by A-weighting at 15 cm from speaker) or 11 kHz (75 dB). For delay conditioning experiments, one CS (CS+) co-terminated with the unconditioned stimulus of footshock (US; 0.65mA, 0.5 s), and the other CS (CS-) was not associated with any event. For trace conditioning experiments, the end of CS+ and US was separated by 15 s of no stimulus. In the acquisition phase, animals were injected with either ketamine (10 mg/kg, s.c.) or saline (10 mL/kg, s.c.), returned to the home cage, and then placed in the conditioning box 15 minutes later. For DREADD experiments, CNO (5 mg/kg, i.p.) or vehicle (10 mg/ml, i.p.) was injected 30 min before trace fear conditioning followed by ketamine or saline 15 min before commencement of conditioning. After 6 minutes of habituation, conditioning started during which 5 CS- and 5 CS+ were presented in a random order with a random inter-trial period of 100 – 110 s. We counter-balanced across subjects the auditory characteristics of CS+ and CS-as well as the ketamine and saline injections. In the recall phase, which occurred about 24 hr later, we tested the animals in a different context. Context was altered using different odors, wall inserts, light, and ambient noise. After 6 minutes of habituation, testing commenced, during which 4 CS-were presented followed by 4 CS+ (with no US) with a random inter-trial period of 100 – 110 s. Motion was quantified from the infrared video data using automated procedures in the Video Freeze software (Med Associates, Inc.). Briefly, according to the publication related to the software^71^, motion was quantified by calculating the sum of pixel-by-pixel value changes across successive video frames. Time periods when the motion values fell below the software-preset threshold for at least 1 s were considered freezing periods. For 35 s after the onset of each CS+ or CS-(i.e., the 20-second long CS presentation plus the 15-second long trace period), we determined the fraction of time in which freezing was detected. We reported the differential value, i.e., the fraction of time spent freezing in response to CS+, subtracted by the fraction of time spent freezing in response to CS-, as measured in the recall phase.

#### Pre-pulse inhibition

To measure sensorimotor gating, we used a commercial system (SR-Lab, San Diego Instruments). An animal would be injected with ketamine (40 mg/kg, s.c.) or saline (10 mL/kg, s.c.) and then placed in a cylinder on a platform with an accelerometer for recording motion, all of which lie inside a soundproof chamber. For DREADD experiments, CNO (5 mg/kg, i.p.) or vehicle (10 mg/ml, i.p.) was given 30 minutes before injection of ketamine or saline and immediate commencement of PPI. The animal was habituated for 5 minutes to background noise (66 dB, calibrated by A-weighting at 25 cm from speaker) and startle stimuli (120 dB noise, 50 ms). For testing, there were 5 types of trials: startle, pre-pulse trials during which the onset of a pre-pulse stimulus (3, 6, or 9 dB noise above background, 50 ms) preceded a startle stimulus by 100 ms, and catch trials with no stimulus. A block consisted of 5 trials, 1 of each type, in a random order with a random inter-trial period of 8 – 23 s. For each animal, we would test for 10 blocks (i.e., 50 trials in total). For DREADD experiments, CNO (5 mg/kg, i.p.) or vehicle (10 mL/kg, i.p.) was injected 30 min before trace fear conditioning followed by ketamine or saline 15 min before commencement of conditioning. The pre-pulse inhibition (PPI) is calculated as follows:

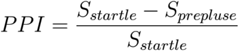

where *S*_startle_ and *S*_prepulse_ were the rectified motion signals recorded by the accelerometer during the 50-ms long startle stimulus averaged across trial repeats, for the startle and prepulse trials respectively.

#### Open-field locomotor activity

To assess locomotion, we placed animals in a conditioning box equipped with an infrared video camera (320 ×8 240 pixels, 30 Hz; Med Associates, Inc.). After approximately 30 minutes of habituation, they were injected with either ketamine (10 mg/kg, s.c.) or saline (10 mL/kg, s.c.) and returned to the box for another 60 minutes. Motion was assessed from the infrared video data quantified using automated procedures in the Video Freeze software (Med Associates, Inc.). To plot the results for **Fig. 5c**, we smoothed the motion signals with a 5-minute long moving window at 1 s steps, for visualization purposes. For statistical comparisons, we used the time-averaged motion for each of the epochs: pre-injection, 5-30 min post-injection and 30-60 min post-injection.

### Statistics

Statistical tests as described in figure captions were done in MATLAB. For the imaging experiments, all planned two-group comparisons were done with parametric t-tests (two-tailed, α = 0.05). For experiments with multiple parameters and conditions, we employed ANOVA tests, with groupings as indicated in the figure caption, followed by post-hoc tests using Tukey-Kramer for statistically significant main or interaction effects. The sampling distribution of the mean was assumed to be normal, but this was not formally tested. To test the robustness of our results, we also ran all the statistical tests in Figs. 1-3 using the two-sample Kolmogorov– Smirnov (K-S) test. This nonparametric test allows us to compare the two cumulative fraction plots (saline versus ketamine) directly without any assumption of normality. None of the conclusions of whether to reject the null hypothesis at α = 0.05 are affected by a change in statistical tests (data not shown). For behavior and electrophysiology experiments, we used non-parametric tests (two-tailed, α = 0.05). In the figures, *P*-values were represented as n.s., not significant; *P* > 0.05; \**P* < 0.05; \*\**P* < 0.01; \*\*\**P* < 0.001. See Figure captions for exact *P*-values.

## Data availability

The datasets that were generated and/or analyzed during the present study are available from the corresponding author upon reasonable request.

## Code availability

Matlab scripts for the present study are available from the corresponding author upon reasonable request.

## Acknowledgements

We thank M. Picciotto for supplying pAAV-CMV-dsRed-pSico-GFP; S. Tomita, M. Higley, and J. Krystal for discussions, I.-J. Kim for use of microscope; V. Phoumthipphavong and M. Rapanelli for technical help. This work was supported by National Institute of Mental Health grants R01MH112750 (A.C.K.) and R21MH110712 (A.C.K.), NARSAD Young Investigator Grant (A.C.K.), Simons Foundation Autism Research Initiative Pilot Award (A.C.K.), Inscopix DECODE Award (A.C.K.), Alzheimer’s Association Research Fellowship AARF-17-504924 (F.A.) and James Hudson Brown-Alexander Brown Coxe Postdoctoral Fellowship (F.A.).

## Author contributions

F.A. and A.C.K. designed the research. F.A. performed surgery, imaging, electrophysiology, behavioral experiments, and histology. D.M.G., S.P., R.S.D. performed molecular cloning, viral packaging, and *in vitro* validation. C.P. assisted with PPI. F.A., K.S. and A.C.K. analyzed all data. F.A. and A.C.K wrote the paper, with input from all other authors.

## Additional information

### Supplementary Information

accompanies this paper.

---

**Supplementary Fig. 1.**
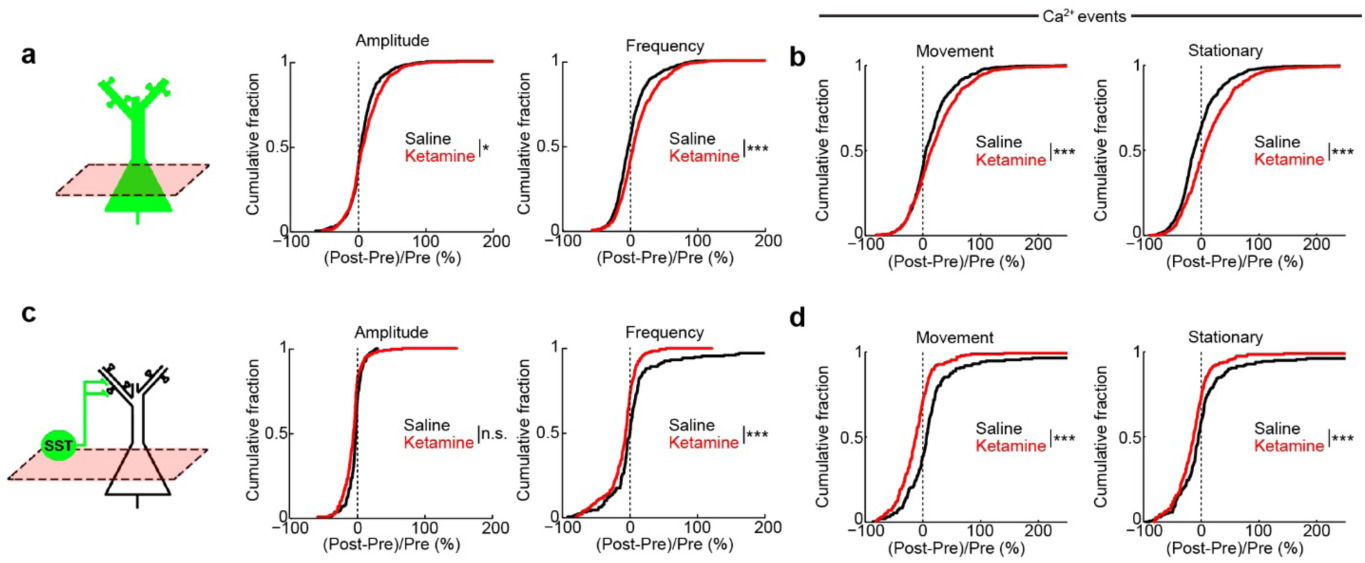
Additional analysis of effects of subanesthetic ketamine on somatic activity of pyramidal neurons and SST interneurons. (a) Left, Schematic of imaging of pyramidal neurons in Cg1/M2. The normalized difference in amplitude (middle) (ketamine: 11 ± 1%, saline: 7 ± 1%, mean ± s.e.m.; *P* = 0.03, two-sample t-test), and frequency (right) of binned calcium events (ketamine: 10 ± 1%; saline: 1 ± 1%; *P* = 3 × 10^-8^, two-sample t-test) for all data regardless of movement. (b) Left, the normalized difference in the rate of spontaneous calcium events of pyramidal neurons during movement (ketamine: 25 ± 2%, saline: 15 ± 2%, mean ± s.e.m.; *P* = 9 × 10^-4^, two-sample t-test). Movement was detected by using a threshold calculated by fitting a two-mean Gaussian to the motion trace and taking two standard deviations above the lower mean associated with stationary periods. Any periods with motion estimate above the threshold were considered movement. Right, same as left for stationary periods (ketamine: 23 ± 2%, saline: 5 ± 2%, mean ± s.e.m.; *P* = 8 × 10^-10^, two-sample t-test). For ketamine, *n* = 613 cells from 5 animals. For saline, *n* = 681 cells from 5 animals. (c) Same as (a) for SST interneurons. Amplitude (ketamine: -4 ± 1%, saline: -1 ± 1%, mean ± s.e.m.; *P* = 0.06, two-sample t-test), and frequency of binned calcium events (ketamine: -10 ± 2%; saline: 12 ± 6%; *P* = 1 × 10^-6^, two-sample t-test). (d) Same as (b) for SST interneurons (movement: ketamine, -12 ± 3%, saline: 19 ± 6%; *P* = 9 × 10^-4^, two-sample t-test; stationary, ketamine, -13 ± 3%, saline: 9 ± 6%; *P* = 9 × 10^-4^, two-sample t-test). For ketamine, *n* = 198 cells from 5 animals. For saline, *n* = 179 cells from 5 animals. * *P* < 0.05; ** *P* < 0.01; *** *P* < 0.001; n.s., not significant

**Supplementary Fig. 2.**
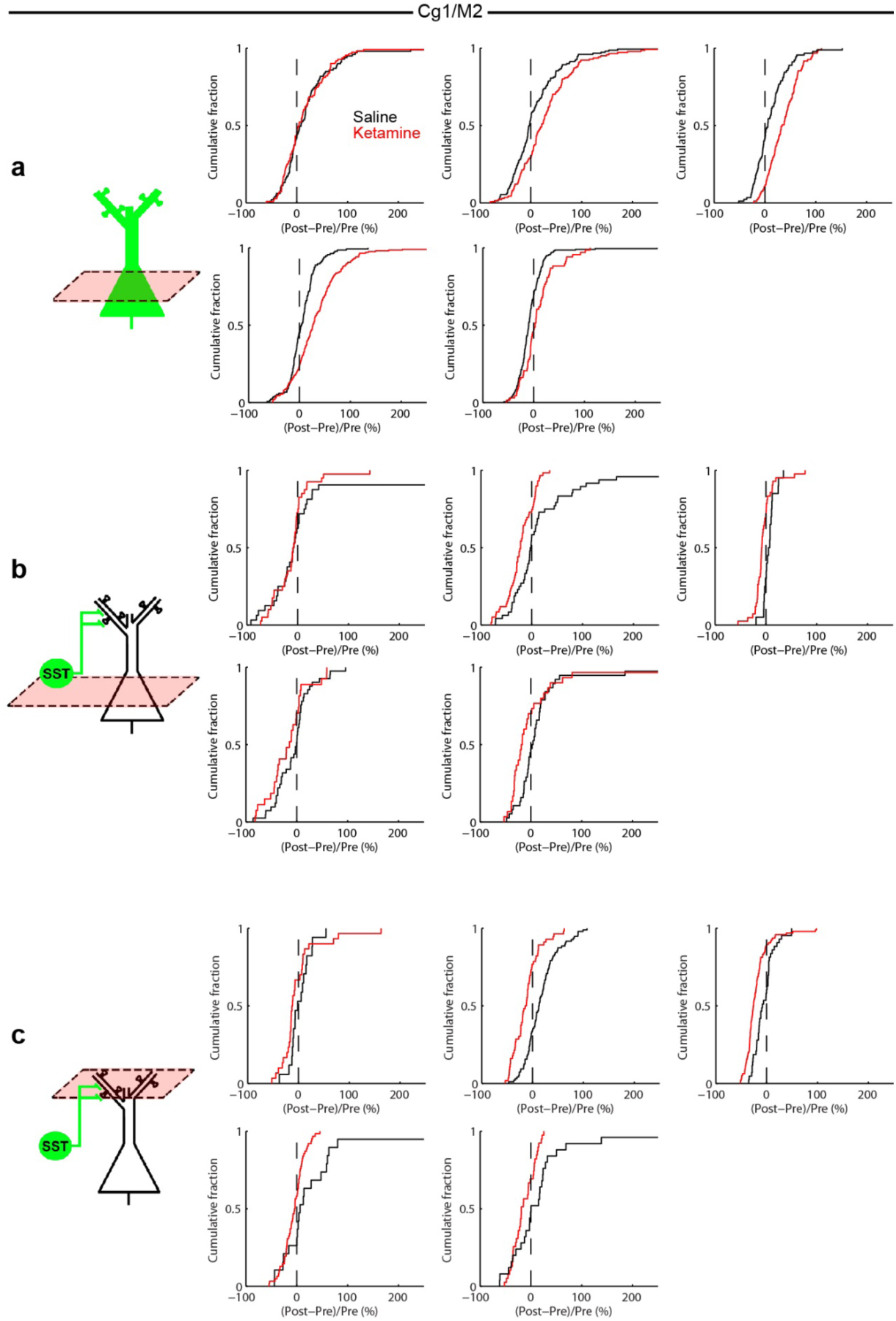
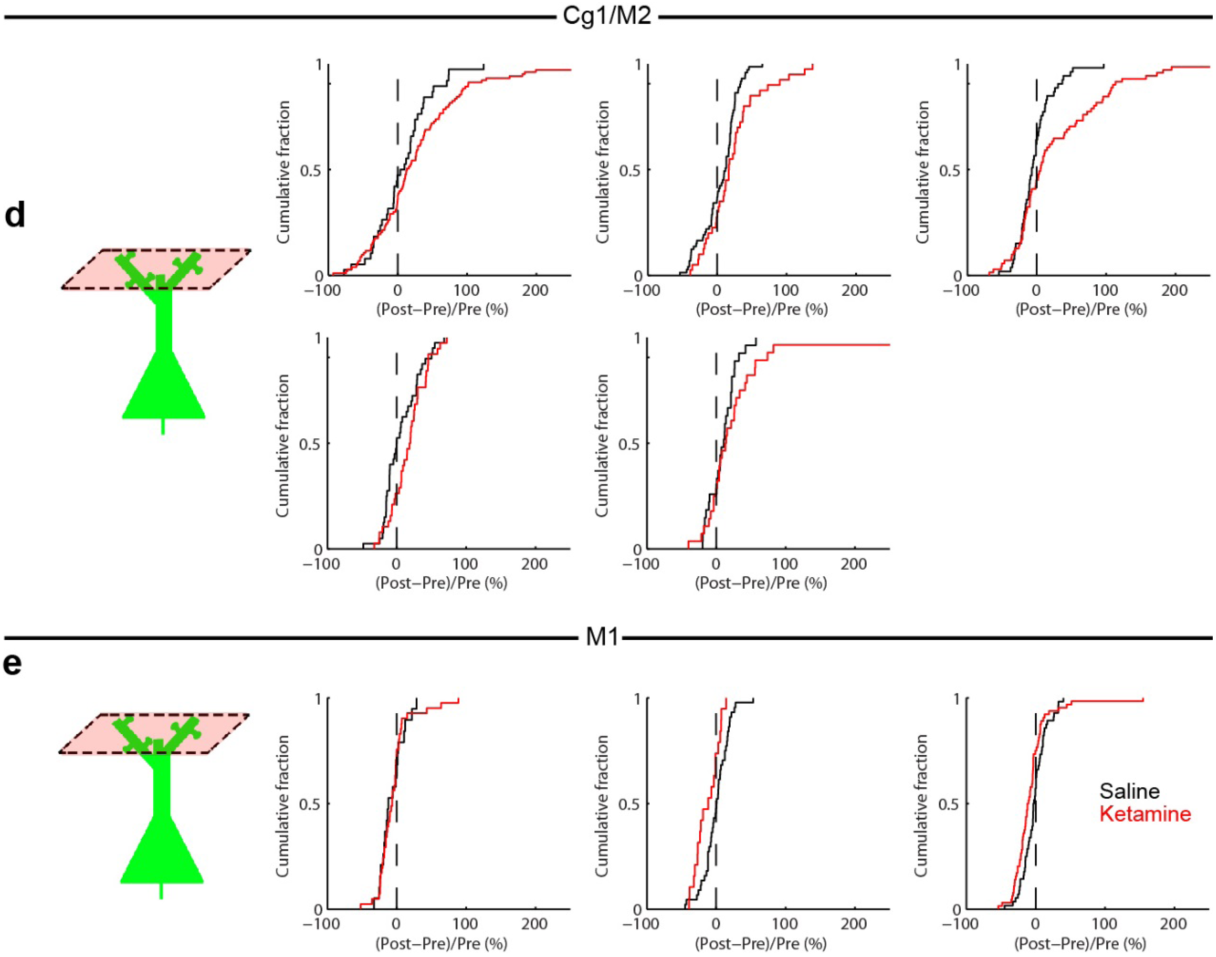
Cumulative fraction plots for individual mice. (a) Schematic of imaging of pyramidal neurons in Cg1/M2. Each plot is the normalized difference (saline, ketamine) in the rate of spontaneous calcium events for an individual mouse. (b) Same as (a) for SST interneuron soma in Cg1/M2. (c) Same as (a) for SST axons in Cg1/M2. (d) Same as (a) for dendritic spines in Cg1/M2. (e) Same as (a) for dendritic spines in M1.

**Supplementary Fig. 3.**
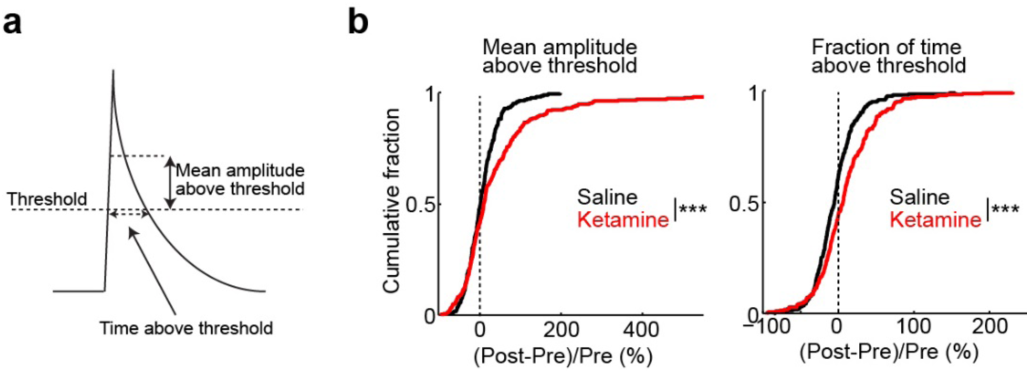
Analyzing the calcium dynamics in dendritic spines with an alternative method. (a) Schematic illustrating an alternative method to detect calcium events from spine fluorescence transients. For each spine, the threshold was defined as 3 times the median absolute deviation of ΔF/F values across all image frames. We identified image frames in which ΔF/F was above the threshold. From this subset of image frames, we determined the mean amplitude above threshold (Δ*F/F*(*t*) minus the threshold), and the time above the threshold (the number of image frames divided by the frame rate). (b) Summary of calcium dynamics for apical dendritic spines in Cg1/M2. Left, normalized difference in the mean amplitude above threshold. Normalized difference was calculated as post-minus pre-injection values normalized by the pre-injection value (ketamine: 58 ± 13%, mean ± s.e.m.; saline: 6 ± 3%; *P* = 5 × 10^-4^, two-sample t-test). Right, normalized difference in the fraction of time of an imaging session spent above threshold (ketamine: 10 ± 2%; saline: -3 ± 2%; *P* = 5 × 10^-5^, two-sample t-test). For saline, *n* = 231 dendritic spines from 5 animals. For ketamine, *n* = 280 dendritic spines from 5 animals. * *P* < 0.05; ** *P* < 0.01; *** *P* < 0.001; n.s., not significant.

**Supplementary Fig. 4.**
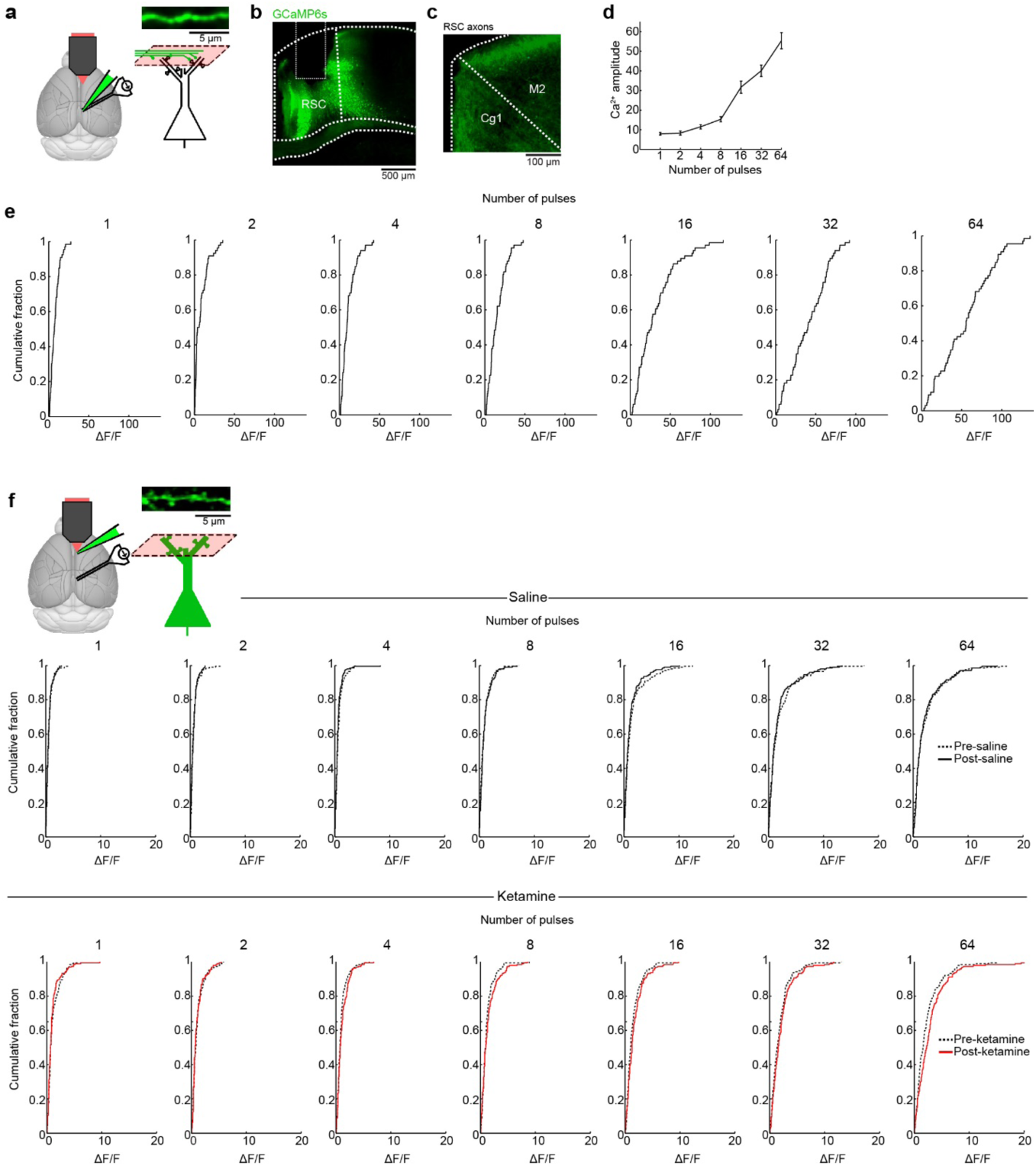
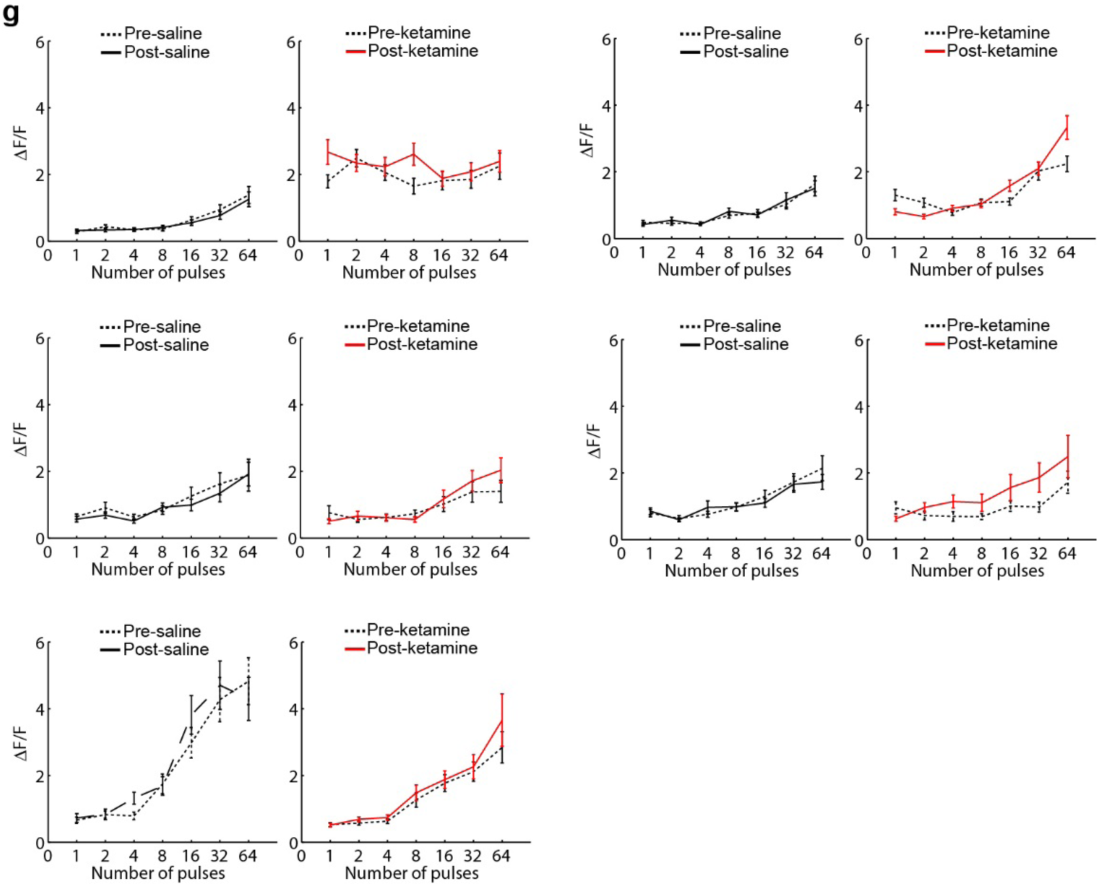
Imaging stimulation-induced calcium responses in RSC axons and spines in Cg1/M2. (a) Schematic of experimental setup and imaging location. A simulation electrode was placed in the retrosplenial cortex (RSC). A virus was injected to mediate GCaMP6s expression in RSC. Axonal calcium responses were imaged in Cg1/M2. (b) Coronal histological section, showing the extent of AAV-mediated expression of GCaMP6s. The white rectangle indicates lesion made by the stimulation electrode. RSC, retrosplenial cortex. (c) Coronal histological section, showing GCaMP6s-expressing axons in the medial prefrontal cortex. Cg1, cingulate cortex. M2, secondary motor cortex. (d) Trial-averaged calcium responses for RSC axonal boutons, as a function of the number of stimulation pulses applied in a trial. Line, mean ± s.e.m. *n* = 66 axonal boutons from 2 animals. (e) Trial-averaged calcium response for RSC axonal boutons, plotted in separate axes for each stimulation level, to show the variability in stimulation-evoked responses of single axonal boutons. These plots are the full distribution of data used to generate panel (D). (f) Trial-averaged calcium response for dendritic spines in Cg1/M2, plotted in separate axes for each stimulation level, to show the variability in stimulation-evoked responses of single dendritic spines. These plots are the full distribution of data used to generate panel Fig. **4**d. (g) Individual mouse data for stimulation-induced calcium responses in spines in Cg1/M2. Five individual mice, with each mouse showing ΔF/F responses pre-saline, post-saline, pre-ketamine and post-ketamine.

**Supplementary Fig. 5.**
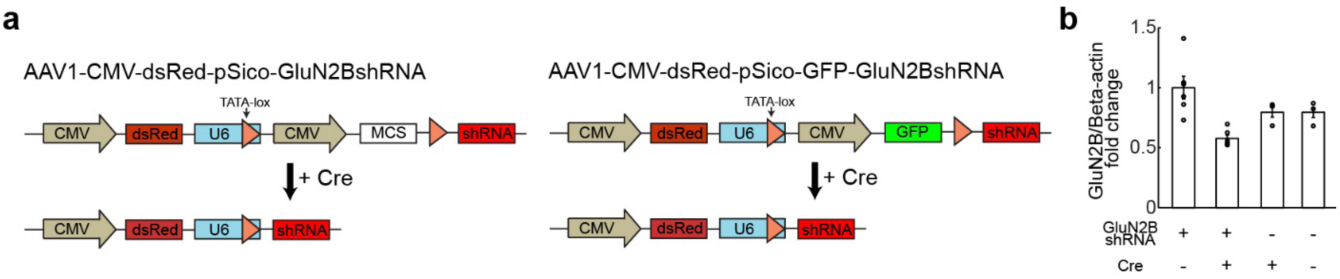
Validation of Cre-dependent knockdown GluN2B receptors. (a) Left, schematic of AAV1-CMV-dsRed-pSico-GluN2BshRNA (standard AAV cassettes not drawn) for Cre-dependent expression of GluN2BshRNA used for combined knockdown and imaging experiments. Right, schematic of AAV1-CMV-dsRed-pSico-GFP-GluN2BshRNA for Cre-dependent expression of GluN2BshRNA used for behavioral experiments. (b) Protein expression levels via Western blot to validate Cre-dependent GluN2B knockdown for AAV1-CMV-dsRed-pSico-GluN2BshRNA. GluN2B signals relative to beta-actin signals for 4 conditions (all values normalized to the GluN2BshRNA, no Cre condition): GluN2BshRNA, no Cre (1.00 ± 0.09, mean ± s.e.m.); GluN2BshRNA, Cre (0.58 ± 0.03), no GluN2BshRNA, Cre (0.80 ± 0.06); no GluN2BshRNA, no Cre (0.79 ± 0.06).

